# Mitochondria-dependent phase separation of disease-relevant proteins drives pathological features of age-related macular degeneration

**DOI:** 10.1101/2020.07.15.201723

**Authors:** Nilsa La Cunza, Li Xuan Tan, Thushara Thamban, Colin J. Germer, Gurugirijha Rathnasamy, Kimberly A. Toops, Aparna Lakkaraju

**Affiliations:** Department of Ophthalmology, School of Medicine, University of California, San Francisco, CA, USA; Pharmaceutical Sciences and Pharmacogenomics Graduate Program, University of California, San Francisco, CA, USA; TurtleTree Labs, Singapore; Zeiss Microscopy, New York; Department of Anatomy, School of Medicine, University of California, San Francisco, CA, USA

## Abstract

Age-related macular degeneration (AMD) damages the retinal pigment epithelium (RPE), the tissue that safeguards photoreceptor health, leading to irreversible vision loss. Polymorphisms in cholesterol and complement genes are implicated in AMD, yet mechanisms linking risk variants to RPE injury remain unclear. We sought to determine how allelic variants in the apolipoprotein E cholesterol transporter modulate RPE homeostasis and function. Using live-cell imaging, we show that inefficient cholesterol transport by the AMD risk-associated ApoE2 increases RPE ceramide, leading to autophagic defects and complement-mediated mitochondrial damage. Mitochondrial injury drives redox state-sensitive cysteine-mediated phase separation of ApoE2, forming biomolecular condensates that could nucleate drusen. The protective ApoE4 isoform lacks these cysteines and is resistant to phase separation and condensate formation. In *Abca4*^-/-^ Stargardt macular degeneration mice, mitochondrial dysfunction induces liquid-liquid phase separation of p62/SQSTM1, a multifunctional protein that regulates autophagy. Drugs that decrease RPE cholesterol or ceramide prevent mitochondrial injury and phase separation in vitro and in vivo. In AMD donor RPE, mitochondrial fragmentation correlates with ApoE and p62 condensates. Our studies demonstrate that major AMD genetic and biological risk pathways converge upon RPE mitochondria, and identify mitochondrial stress-mediated protein phase separation as an important pathogenic mechanism and promising therapeutic target in AMD.

## Introduction

Age-related macular degeneration (AMD), a complex neurodegenerative disease that destroys central high-resolution vision, is the most common cause of permanent blindness in older adults. Global AMD prevalence is projected to exceed 200 million by the year 2040 largely due to lack of disease-modifying therapies for non-neovascular or “dry” AMD, which affects the majority of patients (1, 2). The primary site of injury in AMD is the retinal pigment epithelium (RPE), a monolayer of postmitotic polarized cells that performs numerous functions essential for photoreceptor health and for vision (3, 4). In early AMD, RPE dysfunction is accompanied by the deposition of sub-RPE and sub-retinal lipid-protein aggregates called drusen (5–7). Clinically, RPE abnormalities and drusen location, size, and morphology are strong predictors of AMD progression (1). Polymorphisms in over 50 independent genes involved in regulating complement activation, cholesterol homeostasis, oxidative stress, and extracellular matrix remodeling are associated with AMD. Yet, we have limited insight into the mechanisms by which AMD-associated variants promote RPE injury and drusen formation (7–9).

Of the cholesterol regulatory genes implicated in AMD, the cholesterol transporter *APOE* is especially enigmatic because humans are the only species that express *APOE* allelic variants, which show reversed risk associations between AMD and Alzheimer’s disease. *APOE4* significantly reduces the risk of developing AMD and *APOE2* is associated with increased susceptibility (8, 10–12), whereas the opposite is true for Alzheimer’s disease (13, 14). At least three lines of evidence show that ApoE and cholesterol are major players in AMD pathogenesis: the RPE is the primary biosynthetic source of ApoE in the retina, ApoE secretion by polarized RPE monolayers is increased upon complement exposure (6, 15, 16), and drusen from AMD eyes are rich in cholesterol and ApoE (5). ApoE transports cholesterol in association with ABCA1, another protein implicated in AMD, and

RPE-specific deletion of *ABCA1* in mice causes RPE cholesterol storage and progressive RPE atrophy, photoreceptor loss, and retinal degeneration (17). These studies suggest that isoform-specific functions of ApoE could have a major impact on the RPE homeostasis. However, precisely how *APOE* variants influence RPE health and drusen formation, and how this impacts disease pathogenesis are critical, unresolved questions in AMD biology.

Work from our group has identified complex multilayered interactions between the cholesterol and complement pathways in derailing RPE homeostasis in models of macular degeneration. We have shown that excess RPE cholesterol in the *Abca4*^-/-^ mouse model of Stargardt macular degeneration leads to a secondary accumulation of ceramide, which in turn compromises mechanisms that protect the RPE from complement attack. Sustained complement activation eventually causes mitochondrial injury and metabolic deficits in the RPE (18, 19). Of note, while mitochondrial abnormalities have been documented in AMD patient RPE (20–22), mechanisms connecting loss of mitochondrial integrity with specific features of AMD pathology have yet to be identified.

Here, we sought to investigate how ApoE isoforms regulate cholesterol transport and mitochondrial integrity, and determine how this could contribute to drusen biogenesis. Using high-speed live-cell imaging of polarized adult primary RPE monolayers, *Abca4*^-/-^ mice, and human AMD donor tissues, we show that efficient cholesterol transport in RPE expressing the non-risk ApoE3 or the protective ApoE4 isoforms preserves critical homeostatic functions such as autophagy despite cellular stress. In contrast, ApoE2-expressing RPE accumulate cholesterol, making the RPE susceptible to complement-mediated mitochondrial fragmentation. Our data identify an exciting new mechanism of redox-mediated liquid-liquid phase separation in the RPE. We show that fragmented mitochondria promote ApoE2 phase separation as a result of thiol oxidation of cysteines at 112 and 158, resulting in the formation of biomolecular condensates as potential drusen precursors. The AMD protective ApoE4, with arginines at both positions, is resistant to phase separation, providing a biophysical basis for ApoE risk associations in AMD. RPE mitochondrial injury drives redox state-dependent phase separation of other proteins with reactive cysteines, notably the multifunctional protein p62/SQSTM1. As p62 regulates autophagy, the proteasome, and the antioxidant stress response, this would have far-reaching consequences for RPE health. Drugs that decrease RPE cholesterol or ceramide prevent mitochondrial injury, and limit the formation of biomolecular condensates in *Abca4*^-/-^ mice. Using human tissue as an ex vivo model, we observed significantly more mitochondrial fragmentation and larger ApoE and p62 condensates in AMD donor RPE.

This study integrates genetic and biological risk pathways with poorly understood features of AMD: cholesterol and complement dysregulation, mitochondrial injury, and drusen biogenesis. Our data provide nuanced insight into how a key genetic risk variant mediates crosstalk between lipid and complement pathways to disrupt critical metabolic functions, and identifies mitochondria as a mediator of liquid-liquid phase separation of proteins implicated in AMD. Our studies also suggest that targeting upstream mechanisms that preserve redox homeostasis could be a promising therapeutic approach to limit phase separation and safeguard RPE health.

## Results

### Isoform-specific dynamic ApoE trafficking alleviates pathological cholesterol accumulation in the RPE

In the human *APOE* gene, two polymorphisms (rs7412 and rs429358) result in cysteine/arginine interchanges at positions 112 and 158 in the ApoE protein: ApoE2 has two cysteines, ApoE3 has a cysteine and an arginine, and ApoE4 has two arginines (Figure S1A). The single amino acid changes between ApoE2, E3 and E4 substantially alter protein structure and function, including oligomer formation, and binding to lipids and receptors (23–27). *APOE3* is the most prevalent allele and *APOE2* the least (~79% global allele frequency for *APOE3* compared to ~7% for *APOE2* and 14% for *APOE4)* (28).

To determine how ApoE isoforms modulate steady-state cholesterol levels in healthy and stressed RPE, we first performed high-speed live-cell imaging of primary RPE expressing mCherry-tagged ApoE2, ApoE3 or ApoE4 (Figure S1A). We observed comparable transfection efficiencies and expression of human ApoE fusion proteins measured by immunoblotting and single-cell mCherry fluorescence (Figures S1B and S1C). Four-dimensional analysis of ApoE vesicle trajectories showed that ApoE3 and ApoE4 vesicles exhibited long-range, directed movements (large displacements) compared to ApoE2 (Figures 1A & 1B) likely due to structural and conformational differences between the isoforms. Intermolecular disulfide bonds between cysteines at positions 112 and 158 in ApoE2 (Figure S1A) could induce the formation of multimers that are retained within the cell, as reported for macrophages and adipocytes (29, 30). Additionally, ApoE4 has a closed conformation due to domain interactions via salt bridges between the N-terminal receptor binding domain and the C-terminal lipid-binding domain, whereas ApoE2 adopts the most open conformation and ApoE3 an intermediate conformation (31). Thus, the more compact, dynamic structures of ApoE3 and ApoE4 relative to ApoE2 would enable efficient intracellular trafficking.

**Figure 1.**
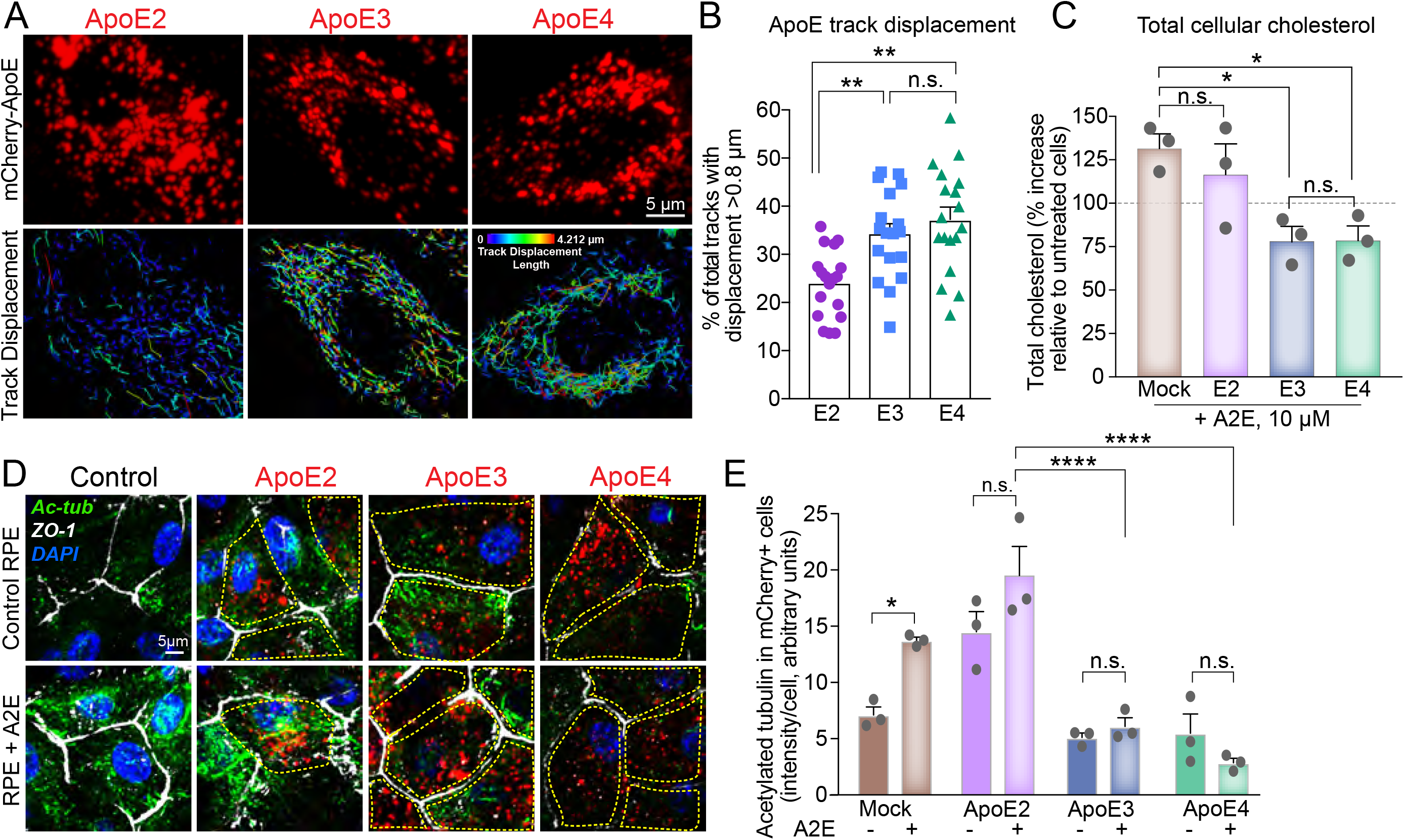
ApoE isoform-specific differences in regulating RPE cholesterol transport and microtubule dynamics. (A) Stills from live imaging of mCherry-tagged human ApoE2, E3, or E4 (top panel, red) in primary RPE cultures. Lower panel: tracks of individual ApoE vesicles. Color bar shows displacement of individual tracks from short (cooler colors) to long (warmer colors), range 0.0 μm to 4.212 μm. (B) Percent of total ApoE tracks with displacement > 0.8 μm in ApoE2, E3 or E4-expressing RPE. Mean ± SEM, n >18 cells per condition from three independent experiments. **, p < 0.005. n.s. - not significant. (C) Endogenous cholesterol content in mock-transfected or ApoE2, E3 or E4-expressing RPE treated with A2E. Mean ± SEM from three independent experiments, 3 replicates/experiment. (D) Representative images and quantitation of acetylated tubulin (green) immunostaining in mock-transfected or ApoE2, E3 or E4-expressing RPE treated or not with A2E. Cell boundaries are demarcated by ZO-1 (white) and nuclei are labeled with DAPI (blue). Cells expressing mCherry-ApoE (red) are outlined in yellow. (E) Quantification of acetylated tubulin intensity in mCherry-expressing cells. Mean ± SEM, n = 10 cells/condition. B, C, and E: One-way ANOVA with Bonferroni post-test. *,*p* < 0.01; **,*p* < 0.001; ****,*p* < 0.0001. n.s. - not significant. See also Figure S1.

Pathological accumulation of lipofuscin bisretinoids, vitamin A metabolites of the visual cycle, is associated with inherited and age-related macular degenerations (32). Because bisretinoids such as A2E cause a secondary accumulation of cholesterol in the RPE (18, 19, 33, 34), we asked how ApoE isoforms would impact the ability of the RPE to deal with excess cholesterol. Biochemical measurements showed that expression of ApoE3 or ApoE4, but not ApoE2, prevented A2E-induced increase in total cell cholesterol (Figure 1C).

Bidirectional lipoprotein secretion by polarized RPE shuttles free and esterified cholesterol into and out of the retina (35). We used filipin (33) and Bodipy 493/503 (36), respectively, to measure free cholesterol and neutral lipids in RPE expressing ApoE2, E3, or E4. Compared to ApoE3- and ApoE4-RPE, ApoE2-RPE had significantly more lipid droplets (Figures S1D and S1E) and free cholesterol (Figure S1F), reflecting their differential trafficking dynamics.

Cell cholesterol modulates organelle biogenesis and trafficking by regulating membrane dynamics and association with microtubules (37). We have shown that lipofuscin-mediated cholesterol storage in the RPE leads to aberrant activation of acid sphingomyelinase (ASMase), the enzyme that hydrolyzes sphingomyelin to ceramide. Increased ceramide results in the accumulation of stable acetylated microtubules, which interferes with autophagosome trafficking, endosome biogenesis, and lysosome localization (18, 19, 33). Immunostaining of ApoE2-, ApoE3-, and ApoE4-expressing RPE for acetylated tubulin showed that expression of either ApoE3 or ApoE4 rescues A2E-mediated increase in tubulin acetylation in the RPE, whereas ApoE2 does not (Figures 1D & 1E).

Taken together, these studies indicate that ApoE2 exhibits significant defects in intracellular trafficking and cholesterol transport, likely due to cysteines at 112 and 158, which promote intracellular retention (29-31, 38). Further, dynamic long-range trafficking of ApoE3 and ApoE4 is essential for maintaining cholesterol homeostasis and microtubule-based trafficking in the RPE.

### ApoE2 aggravates autophagic defects induced by lipofuscin bisretinoids

To understand how ApoE isoform-specific cholesterol transport impacts critical RPE functions, we investigated autophagy, an evolutionarily conserved mechanism responsible for degrading and recycling cellular debris. Lipofuscin-induced cholesterol accumulation and disruption of microtubule-based transport interfere with autophagy in *Abca4*^-/-^ mice and in primary RPE cultures with A2E (33). These autophagic defects manifest as decreased autophagosome biogenesis (decreased LC3B lipidation) and trafficking and impaired autophagic flux.

We performed high-speed live-cell imaging of EGFP-LC3-labeled autophagosomes in ApoE2, ApoE3, or ApoE4-expressing primary RPE (Figures 2A and S2A; Supplementary Movies S1-S8). As expected, A2E decreased autophagosome biogenesis in mock-transfected RPE (33). Expression of ApoE3 or ApoE4, but not ApoE2, increased autophagosome numbers in RPE with A2E comparable to control levels (Figure 2B). Analysis of LC3 trafficking showed that autophagosomes in RPE with A2E had predominantly shorter tracks, with significantly lower mean speeds and displacements (Mock-transfected control vs mock-A2E, Figures 2C-E; Supplementary Movies S1 and S2). RPE expressing ApoE3 or ApoE4 were resistant to A2E-induced disruption of autophagosome trafficking, whereas ApoE2 expression was ineffective (Figures 2C-E and S2B-G; Supplementary Movies S3-S8). These data suggest that improved microtubule dynamics in ApoE3- and ApoE4-RPE are essential for stressed RPE to maintain efficient autophagy.

**Figure 2.**
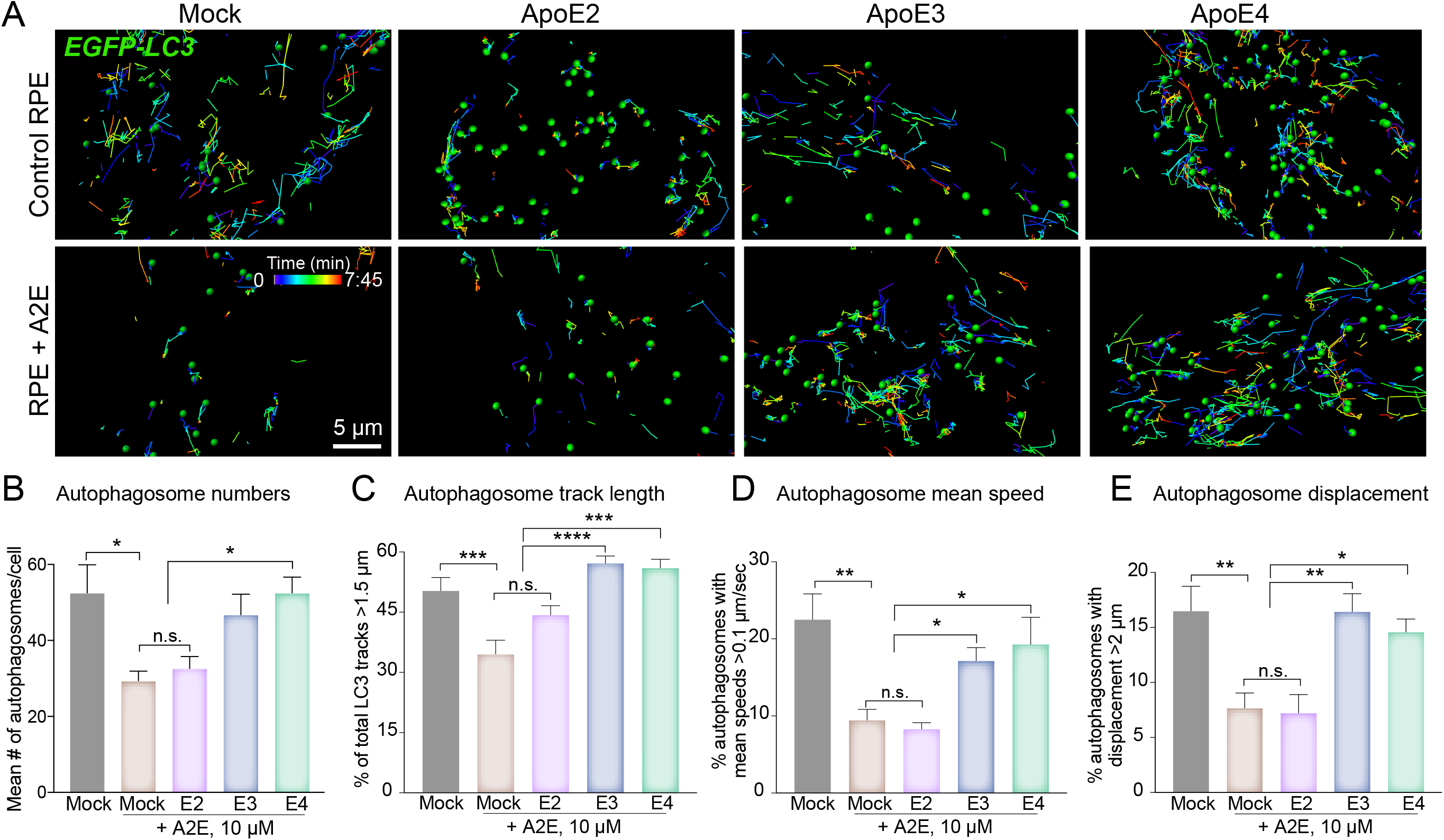
ApoE2 aggravates autophagic defects induced by lipofuscin bisretinoids. (A) Stills from spots and tracks analyses of live imaging of EGFP-LC3 autophagosome trafficking in mock-transfected or mCherry-ApoE2, E3 or E4-expressing RPE, treated or not with A2E. (B) Average number of EGFP-LC3 autophagosomes per cell. (C) Percent of total EGFP-LC3 tracks longer than 1.5 μm. (D) Percent of autophagosomes with mean speeds >0.1 μm/sec. (E) Percent of autophagosomes with displacement > 2 μm in mock-transfected or ApoE2, E3 or E4-expressing RPE, treated or not with A2E. Mean ± SEM, n >13 cells per condition. *, p < 0.05, **, p < 0.005, ***, p < 0.001 and ****, p < 0.0001. n.s. not significant. B to E: One-way ANOVA with Bonferroni post-test. See also Figure S2 and Movies S1-S8.

Decreased autophagosome biogenesis and autophagic flux have been reported in AMD donor RPE (20, 39). Our data identify RPE cholesterol as an important determinant of autophagy, and suggest that protective variants of AMD-associated cholesterol pathway genes could modulate RPE health by preventing autophagic deficits.

### ApoE2 increases RPE susceptibility to complement-mediated mitochondrial injury

Abnormal activation of the alternative complement pathway is strongly associated with AMD. The RPE, which forms the outer blood-retinal barrier, is the first line of defense against uncontrolled complement activation. We have reported that in models of macular degeneration, acetylated microtubules prevent recycling of the complement-regulatory protein CD59 to the plasma membrane and inhibit membrane repair by lysosome exocytosis. This enables uncontrolled assembly of the C5b-9 terminal membrane attack complex (MAC) pore on the cell membrane, and the resulting increase in intracellular calcium causes mitochondrial fragmentation and oxidative stress in the RPE (19). To understand how AMD-associated genes (e.g., ApoE) interact with biological risks (e.g., lipofuscin, cholesterol, complement) to promote mitochondrial injury, we performed live-cell imaging of RPE mitochondrial networks before and after exposure to normal human serum (NHS) as a source of complement (19). Reconstruction of mitochondrial volumes showed highly integrated mitochondria in mock-transfected and ApoE3- or E4-expressing RPE, in agreement with our previous finding that healthy RPE are resistant to complement attack. In contrast, mitochondria in ApoE2-expressing RPE were fragmented in response to NHS (Figures 3A & 3B), indicating that complement-regulatory mechanisms are impaired in ApoE2-RPE, likely due to increased tubulin acetylation. As expected, A2E rendered mock-transfected RPE vulnerable to NHS. In contrast to ApoE2-RPE, expression of ApoE3 or ApoE4 suppressed mitochondrial fragmentation in RPE with A2E after NHS exposure (Figures 3B & 3C), as a consequence of better cholesterol homeostasis, which decreases RPE ceramide, preserves dynamic microtubule trafficking and thereby efficient mechanisms to combat complement activation.

**Figure 3.**
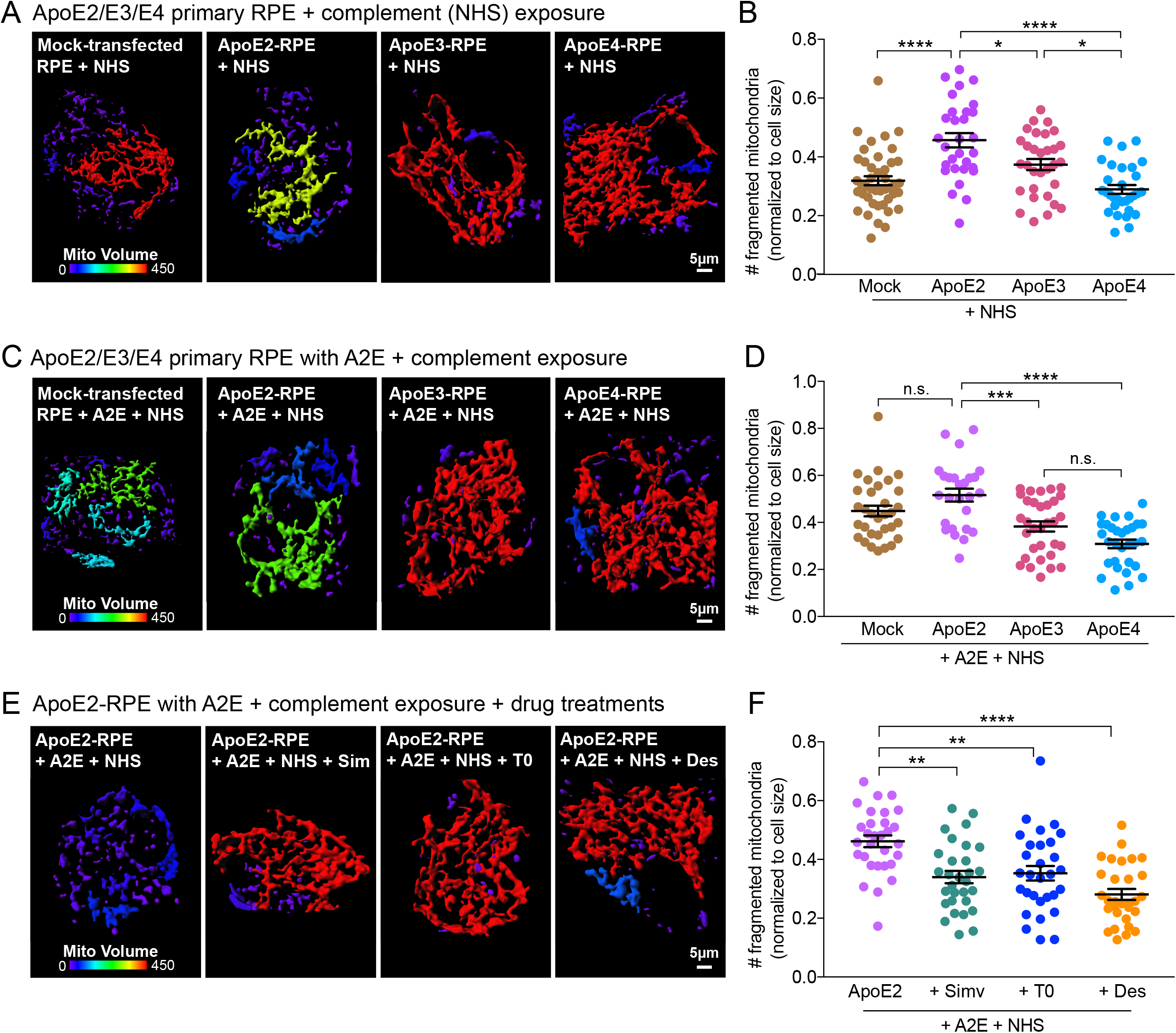
ApoE2 exacerbates complement-induced mitochondrial fragmentation in the RPE. (A) 3D reconstruction of mitochondrial volumes from live imaging of MitoTracker in mock-transfected or mCherry-ApoE2, E3, or E4-expressing RPE exposed to 10% NHS to induce complement attack. Color bar: cooler colors indicate increasing mitochondrial fragmentation. (B) Number of fragmented mitochondria after NHS exposure. (C) Mitochondrial volumes as in (A) in RPE treated with A2E prior to NHS exposure. (D) Quantification of mitochondrial fragments as in (B). (E) Mitochondrial volumes as in (A) in ApoE2-expressing RPE with A2E exposed to NHS and treated with Simvastatin (5 μM, 16 h), T0901317 (1 μM, 16 h) or desipramine (10 μM, 3 h) prior to imaging. (F) Quantification of mitochondrial fragments as in (B). Mean ± SEM, *, p < 0.05; **, p < 0.005; ***, p < 0.0005; ****, p < 0.0001. n = 30 cells (A), 26 cells (B) and 30 cells (C) per condition; One-way ANOVA with Bonferroni post-test; See also Figure S3.

If complement-mediated mitochondrial fragmentation is a result of excess cholesterol activating ASMase, which in turn increases ceramide to induce tubulin acetylation, then drugs that decrease cholesterol or inhibit ASMase (19, 33) should maintain RPE mitochondrial integrity after complement attack. In support of this hypothesis, inhibiting cholesterol biosynthesis with the lipophilic statin simvastatin, increasing cholesterol efflux with the liver X receptor (LXR) agonist T0901317, or inhibiting ASMase with desipramine all decreased complement-induced mitochondrial damage in both mock-transfected (Figures S3A and S3B) and ApoE2-expressing RPE with A2E after exposure to NHS (Figures 3E and 3F). Taken together, these data demonstrate that genetic variants or pharmacological approaches that limit excess cholesterol safeguard the RPE from complement attack and protect mitochondrial integrity.

### ApoE exhibits liquid-like behavior and undergoes phase separation in the RPE

Accumulation of ApoE- and cholesterol-rich aggregates called drusen in the sub-retinal and sub-RPE space is a hallmark of AMD. Clinically, drusen size, shape, and abundance correlate with AMD progression. While studies suggest that the RPE is the minimal requirement for drusen formation and complement exposure increases ApoE secretion by RPE cells in culture (16, 40, 41), molecular mechanisms responsible for drusen biogenesis are unknown. To determine how AMD-associated stressors can nucleate drusen within the RPE, we hypothesized that mitochondrial dysfunction in the RPE could induce ApoE to undergo liquid-liquid phase separation to form biomolecular condensates (42–45). Proteins that undergo phase separation are characterized by intrinsically disordered regions (IDRs) that enable them to engage in low-affinity interactions with one another (46). As ApoE isoforms have IDRs (Figure S4A-C) (47), it is plausible that aberrant phase transitions could drive ApoE condensate formation in the RPE as precursors to drusen.

Phase-separated liquid-like condensates (i) are roughly spherical shape; (ii) fuse with one another to minimize surface tension; and (iii) dynamically respond to changes in the subcellular environment (46, 48–51). Live-cell imaging showed that ApoE forms spherical structures with average sphericity ~0.9 (Figures 4A and 4B) that readily fuse and relax (Figure 4C), suggesting that ApoE2, ApoE3, and ApoE4 all segregate into liquid-like assemblies within the RPE. Although the three isoforms had similar disorder tendencies when analyzed by Meta-predictors such as DisMeta (49) and other commonly used disorder predictors (Figures S4A-C), there were significantly more ApoE2 condensates with larger volumes (> 0.6 μm^3^), compared to either ApoE3 or ApoE4 condensates (Figures 4D and S4D-F). To confirm that ApoE2 forms liquid-like condensates, we treated ApoE2-expressing RPE for two minutes with 1,6-hexanediol, an aliphatic alcohol known to disrupt weak hydrophobic interactions that anchor biomolecular condensates (46, 48, 49). Consistent with our hypothesis of ApoE2 phase separation, we observed that 1,6-hexanediol treatment rapidly decreased both the number and volume of ApoE2 condensates in the RPE (Figures 4E and S4G). These data show that ApoE undergoes phase separation in an isoform-specific manner in the RPE.

**Figure 4.**
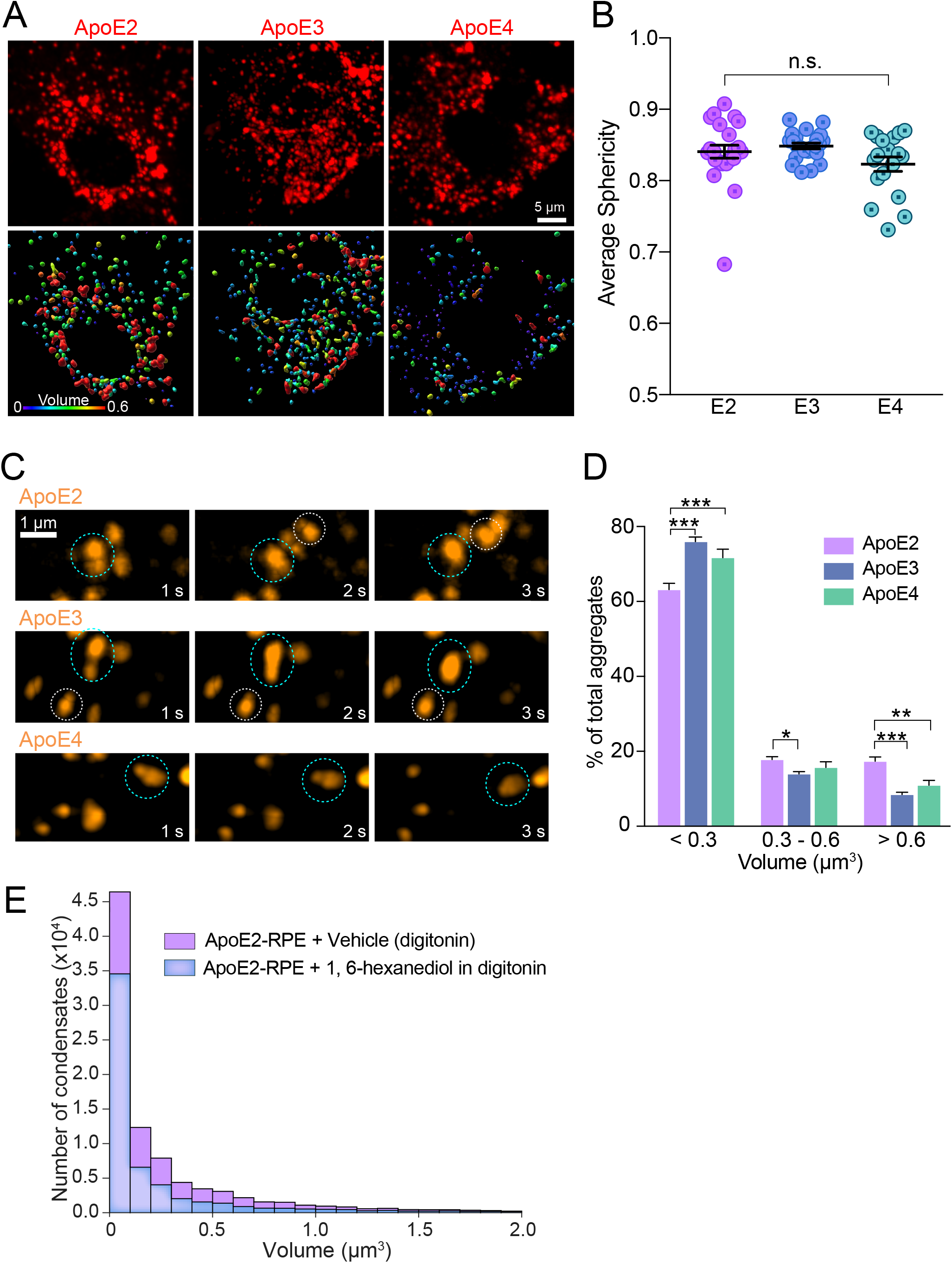
ApoE exhibits liquid-like behavior in the RPE. (A) Representative stills (top panel) and 3D volume reconstructions (lower panel) from live imaging of intracellular mCherry-tagged ApoE2, E3 or E4 in the RPE. Warmer colors indicate larger volumes. (B) Average sphericity of ApoE condensates in RPE expressing ApoE2, E3 or E4. Mean ± SEM, n=17-23 cells per condition. n.s. - not significant. (C) Time-lapse imaging shows that ApoE condensates exhibit liquid behavior by fusing with each other (dotted circles). (D) Frequency distribution of condensate volumes in ApoE2, E3 or E4 expressing RPE. (E) Number and volume of condensates in ApoE2-expressing RPE treated with 1,6-hexanediol, which disrupts weak hydrophobic interactions (blue bars) or with vehicle (digitonin, magenta bars) alone. See also Figure S4. Mean ± SEM, *, p < 0.05; **, p < 0.005; ***, p < 0.0001. n = 15-29 cells per condition; One-way (B) or Two-way (D) ANOVA with Bonferroni posttest.

### Mitochondrial injury drives redox-mediated phase separation of ApoE2 to form biomolecular condensates in the RPE

As liquid-liquid phase separation is known to be exquisitely sensitive to intracellular environment (52), we next asked whether declining metabolic activity in ApoE2-RPE due to loss of mitochondrial function could increase its propensity for phase separation. We analyzed condensate volumes in ApoE2-, ApoE3- and ApoE4-expressing RPE under conditions known to cause cellular stress and mitochondrial damage (i.e., exposure of RPE with A2E to complement). Our data show that whereas A2E alone did not noticeably impact condensate numbers or volumes, exposure of ApoE2-RPE with A2E to NHS significantly increased the number of large (> 0.6 μm^3^) condensates (Figure 5A), consistent with our mitochondrial fragmentation data (Figures 3A-3D). We also observed a small but significant increase in large condensates in ApoE3-RPE exposed to A2E and NHS, whereas ApoE4-RPE were completely resistant to this stress-induced phase separation (Figure 5A). To better understand the relationship between ApoE condensates and mitochondrial function, we used IUPRED2A (53) to predict redox-state dependent transitions in ApoE isoforms. This modeling showed that ApoE2 undergoes more order-disorder phase transitions under oxidative stress compared to ApoE3 or ApoE4 (Figures 5B and S4H-J). ApoE2 has cysteine residues at positions 112 and 158, which can undergo reversible thiol oxidation in response to the intracellular redox environment. Pertinently, these thiol modifications enable ApoE2 to form disulfide-linked homodimers and higher-order oligomers (54–56), which would explain the increase in ApoE2 condensate volumes. In agreement with this model, ApoE3, which has a cysteine at 112 and an arginine at 158, is predicted to have fewer phase transitions under oxidative stress; however, thiol modification at 112 could presumably explain the small increase in condensate formation we observed in Figure 4F. In contrast, ApoE4 has arginines at both 112 and 158, and is therefore impervious to redox-state dependent phase separation and condensate formation (Figures 5B and S4J).

**Figure 5.**
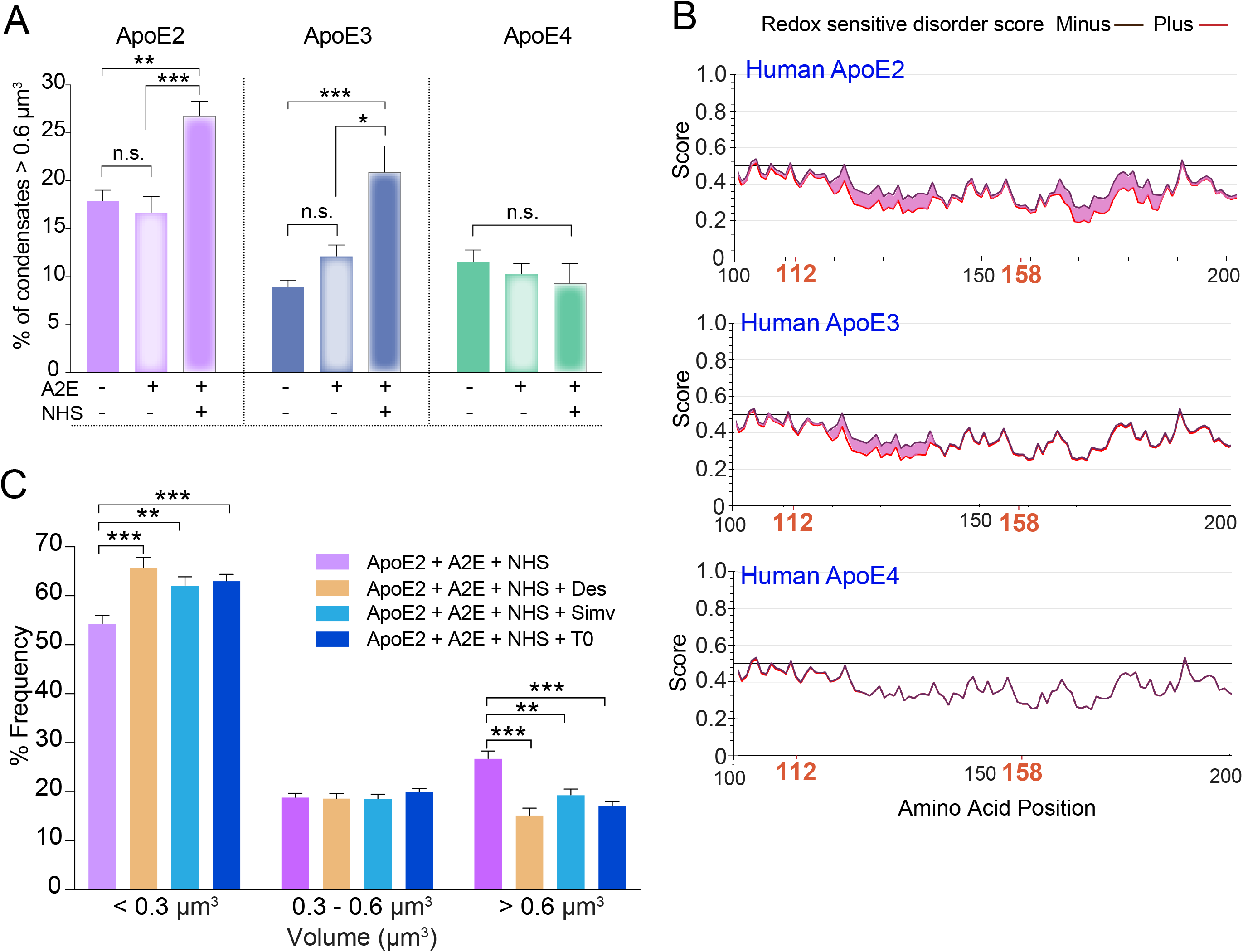
Mitochondrial injury drives redox-mediated phase separation of ApoE2 and ApoE3. (A) Percent of ApoE condensates with volumes greater than 0.6 μm3 in RPE expressing ApoE2, E3 or E4 and treated or not with A2E and NHS. (B) IUPRED2A disorder plots for human ApoE2, ApoE3, and ApoE4. Redox-state dependent order-disorder transitions are depicted by the pink shaded area. Note increased disorder corresponding to cysteines at 112 and 158 in ApoE2 and at 112 in ApoE3. (C) Frequency distribution of condensate volumes in ApoE2-expressing RPE with A2E exposed to NHS and treated with Simvastatin (5 μM, 16 h), T0901317 (1 μM, 16 h) or desipramine (10 μM, 3 h). Mean ± SEM, *, p < 0.05; **, p < 0.005; ***, p < 0.0001. n = 15-29 cells per condition; One-way (A) or Two-way (C) ANOVA with Bonferroni post-test. See also Figure S4.

Finally, to confirm the role of mitochondrial stress in driving ApoE2 phase separation, we asked if drugs that prevent mitochondrial injury after complement attack limit condensate formation. Treatment with either desipramine, simvastatin, or T0901317 significantly decreased the fraction of large ApoE2 condensates in RPE with A2E exposed to complement (Figure 5C). Collectively, these findings identify aberrant phase separation as a powerful and novel mechanism that could nucleate intracellular ApoE2 (and, presumably, ApoE3) condensates within the RPE, especially under conditions of declining mitochondrial health. Further, our data show that pharmacological approaches that safeguard RPE mitochondrial function limit the formation of ApoE condensates.

### Safeguarding mitochondrial health prevents pathological protein phase transitions in the RPE in vivo

Studying the role of mitochondrial stress in ApoE phase transitions in mice is hampered by the fact that mice express a single form of ApoE. Mice with targeted replacement of human *APOE2, APOE3* and *APOE4* (APOE-TR mice) have been widely used to study the role of *APOE4* in Alzheimer’s disease. However, ~100% of mice expressing human *APOE2* develop type III hyperlipoproteinemia, unlike humans, where only 10% of those with the *E2/E2* genotype develop the disease (57). As *APOE2* mice have increased plasma levels of chylomicrons and VLDL remnants, this will likely influence lipoprotein and cholesterol levels in the RPE and retina, which could lead to confounding results in our experiments. These issues could likely explain the differing retinal phenotypes observed in studies using the APOE-TR mice (58, 59).

To avoid these issues, we decided to investigate redox-mediated phase transitions in the pigmented *Abca4*^-/-^ mouse model of Stargardt inherited macular degeneration, which recapitulates many features of early RPE dysfunction observed in AMD including cholesterol and ceramide accumulation, increased microtubule acetylation, and complement activation (18, 19, 32, 33). Pertinent to this study, we previously reported that lipofuscin-induced cholesterol and ceramide accumulation in *Abca4*^-/-^ mice makes the RPE susceptible to complement-mediated mitochondrial injury as evidenced by loss of optic atrophy 1 (OPA1), which mediates inner mitochondrial membrane fusion) (19). 3D reconstructions of TOM20-stained mitochondria in mouse RPE flatmounts showed a significant decrease in mitochondrial volumes, indicative of mitochondrial fragmentation, in 6-month-old *Abca4*^-/-^ mice compared to age-matched wildtypes (Figures 6A & 6B). Mice express a single form of ApoE with arginines at positions 112 and 158, but without the arginine at residue 61 essential for the domain interactions seen with human ApoE4 (60). Based on the lack of oxidizable cysteines at residues 112 and 158, we hypothesized that mouse ApoE would be resistant to redox state-related phase transitions seen in human ApoE2 and ApoE3. This was confirmed by IUPRED2A modeling (Figure S4K) and immunofluorescence staining for ApoE in mouse RPE flatmounts showed no difference in the number or volume of ApoE puncta between wildtype and *Abca4*^-/-^ mice (Figure 6C).

**Figure 6.**
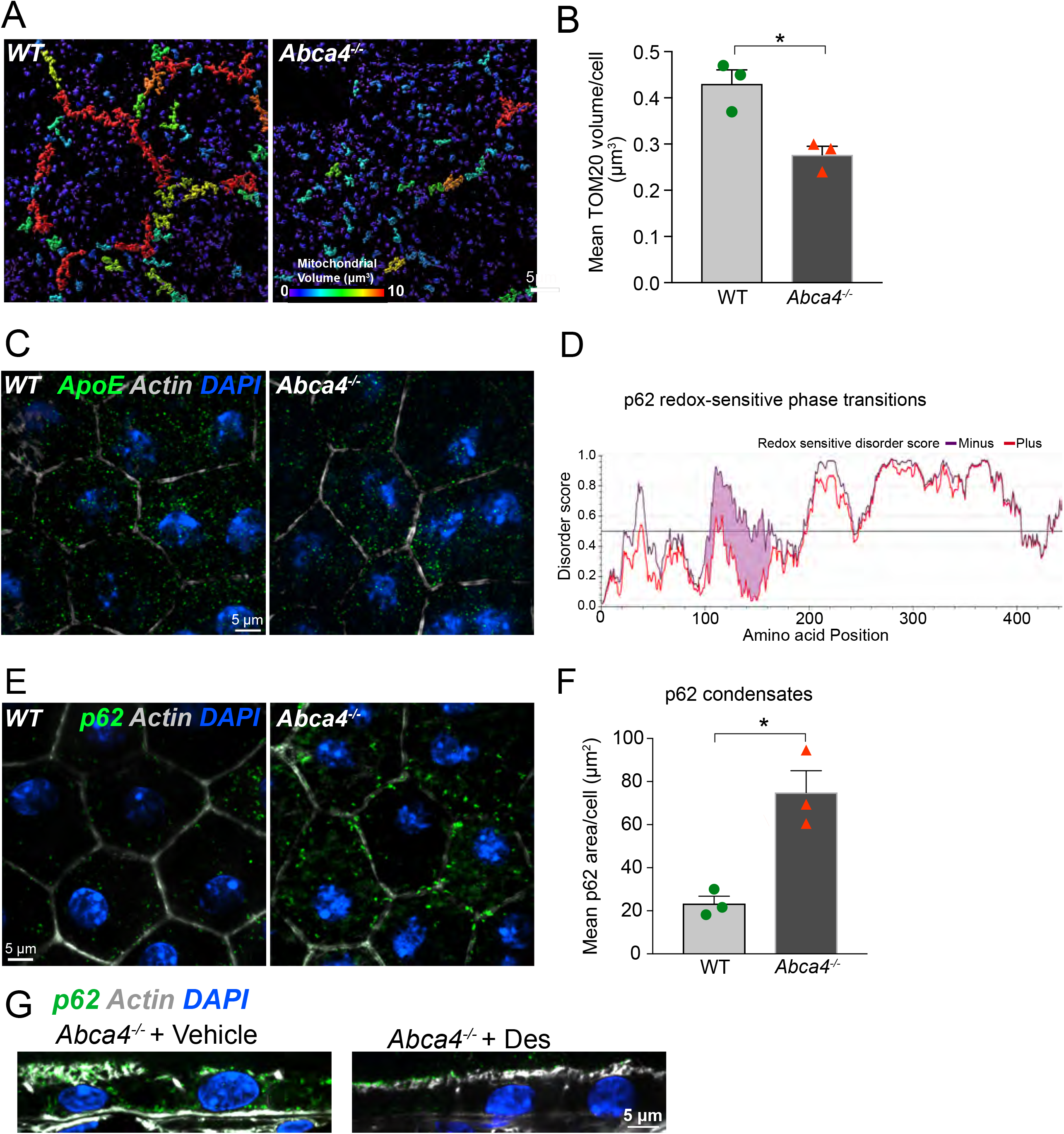
Mitochondrial injury and redox-sensitive protein phase separation in *Abca4*^-/-^ mouse RPE. (A) Volume reconstructions of TOM20-stained mitochondria in wild type and Abca4-/- mouse RPE flatmounts. Color bar - warmer colors indicate larger volumes. (B) ApoE immunostaining (green) in wild type and Abca4-/- mouse RPE flatmounts. (C) IUPRED2 redox-driven disorder predictions for mouse p62. (D) p62 condensates (green) in wildtype and Abca4-/- RPE. (E) p62 (green) in RPE in Abca4-/- mice treated with vehicle or desipramine. B, D, and E: Nuclei - DAPI (blue); Actin - Phalloidin (grey). Mean ± SEM, *, p < 0.05, n = 3 mice per genotype.

We next asked whether our model of mitochondrial redox state-mediated phase separations would be applicable to other proteins with reactive cysteines. We focused on the multifunctional protein p62/SQSTM1, which regulates autophagy, proteasomal clearance, and NRF2-mediated antioxidant response, and has been implicated in AMD pathogenesis (20, 61, 62). IUPRED2A modeling predicted that p62 is highly susceptible to redox state-driven phase transitions because of cysteines at 105 and 113 (63) (Figure 6D). Immunostaining of mouse RPE flatmounts showed numerous large p62 condensates in *Abca4*^-/-^ mice as a consequence of mitochondrial fragmentation compared to age-matched wildtypes (Figures 6E & 6F). Mean areas of p62 aggregates were 23.29 ± 4.94 μm^2^ in wildtype vs 74.8 ± 14.45 μm^2^ in *Abca4*^-/-^ RPE. As further confirmation of the link between mitochondrial health and protein phase separation, a one-month treatment with desipramine, which prevents complement-mediated mitochondrial injury in the RPE, decreased p62 condensates in *Abca4*^-/-^ mice RPE (Figure 6G).

These data demonstrate that mitochondria regulate redox state-mediated abnormal liquidliquid phase separation of proteins with reactive cysteines, driving the formation of intracellular biomolecular condensates in the RPE in vivo. p62 is an autophagy receptor that ferries ubiquitinated cargo to autophagosomes, and abnormal sequestration of p62 in condensates could underlie the autophagic defects reported in *Abca4*^-/-^ mice RPE (33) and in AMD donor RPE (20). Our studies also demonstrate that condensate formation is reversible, and identify pharmacological approaches that limit p62 condensate formation by preserving RPE mitochondrial health in the *Abca4*^-/-^ Stargardt mouse model.

### Mitochondrial fragmentation correlates with ApoE and p62 aggregates in AMD donor RPE

Building upon our in vivo data, we asked if the mechanism linking increased RPE mitochondrial dysfunction to the formation of biomolecular condensates of AMD-relevant proteins would hold in human donors with AMD. We genotyped retinal tissue from unaffected donors and donors with non-neovascular AMD for AMD-associated genes. Both control and AMD donors harbored a mixture of risk and non-risk alleles (Supplementary Tables S2 and S3) reflecting the genetic complexity of AMD, where many individuals with risk variants do not develop the disease and vice versa (8). Volume renderings of TOM20 immunostaining in macular retinal cryosections showed significantly lower mitochondrial volumes (indicative of fragmentation) in all AMD donors compared to unaffected controls (Figures 7A & 7B).

**Figure 7.**
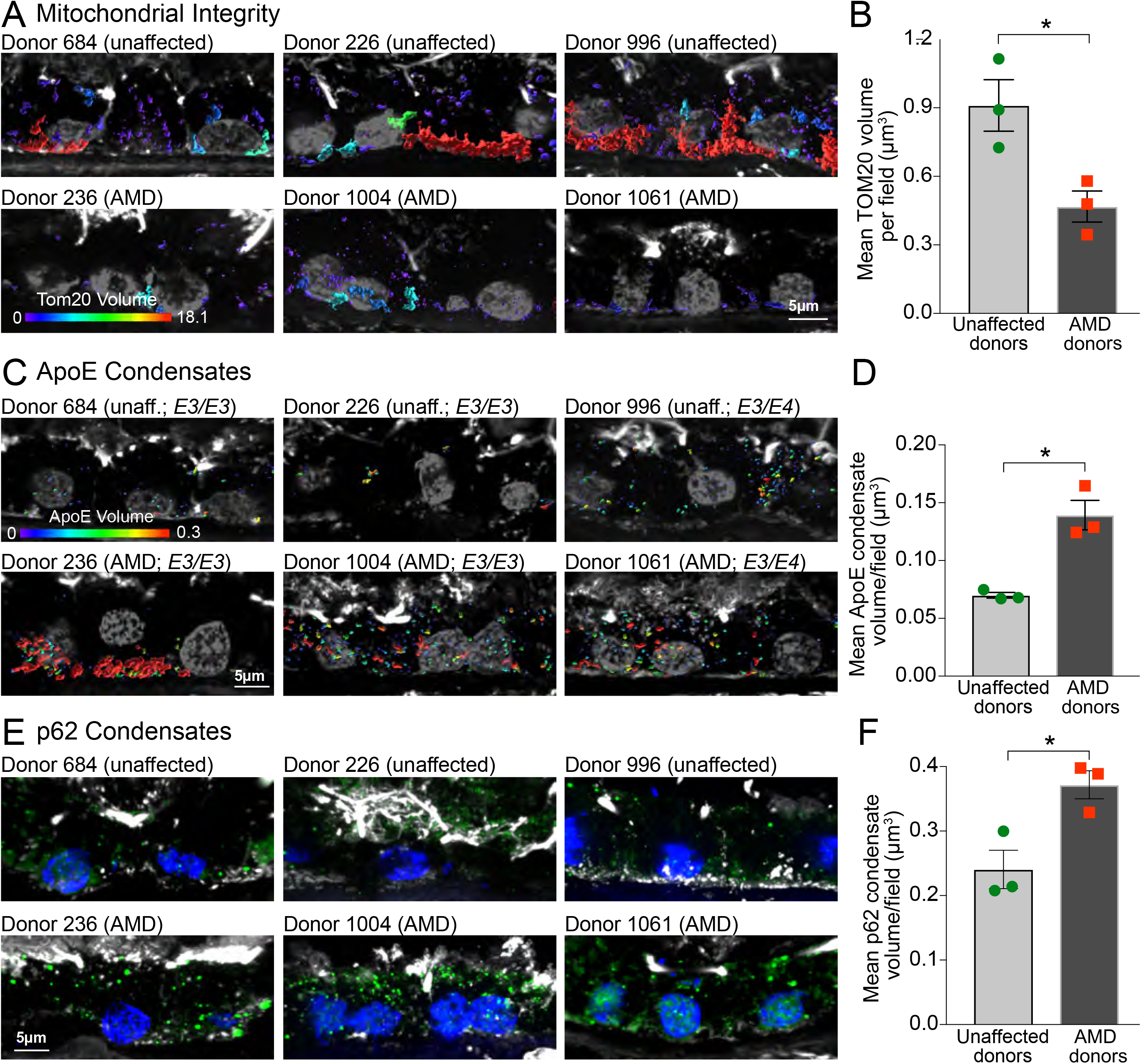
RPE mitochondrial injury correlates with ApoE and p62 condensates in AMD donors. (A) 3D volume reconstructions and (B) quantification of TOM20-labeled mitochondrial volumes in macular RPE cryosections from unaffected donors and donors with AMD. (C) 3D volume reconstructions and (D) quantification of ApoE condensates in human donor RPE. Color bars in (A) and (C): warmer colors indicate larger volumes. (E) p62 immunostaining (green) and (F) quantification of p62 condensates in human donors. In A, C, and E: phalloidin is in white and nuclei are labeled with DAPI (grey in A and C, blue in E). B, D, and F: Data are presented as Mean ± SEM, *, p < 0.05; 3 donors per group. See also Figure S5 and Tables S2 and S3.

Analysis of ApoE (Figures 7C & 7D) and p62 (Figures 7E and 7F) by immunostaining and volume reconstructions showed significantly larger ApoE and p62 condensates in AMD donor RPE compared to unaffected donors. *APOE* allele distribution in both unaffected and AMD donors was identical (two *E3/E3* donors and one *E3/E4* donor each, Table S2). The increased volume of ApoE aggregates in AMD donors with at least one *E3* allele correlates with the mitochondrial fragmentation in these donors, and agrees with data in Figure 5 showing that ApoE3 also undergoes redox-mediated phase separation due to the cysteine at residue 112.

These data, together with our live-cell imaging and mouse studies, provide strong support for a model where mitochondrial injury enables phase separation of AMD-relevant proteins with reactive cysteines, leading to the formation of biomolecular condensates.

## Discussion

Deciphering the roles of AMD-associated genes in the disease process has been challenging because of limited insight into how risk variants modulate RPE homeostasis and contribute to specific features of AMD pathology (9, 64, 65). Here, using in vitro, in vivo, and ex vivo models, we show that differential cholesterol transport mediated by ApoE isoforms (ApoE2≪ApoE3=ApoE4) regulates RPE susceptibility to complement attack. Further, this crosstalk between dysregulated complement and cholesterol pathways converges upon RPE mitochondria, leading to redox-mediated phase separation of ApoE2, the AMD risk isoform. These studies help explain the risk associations of *APOE* allelic variants in AMD and demonstrate how these alleles modulate drusen biogenesis, a hallmark of AMD progression. We show that upstream therapeutic strategies that restore lipid homeostasis prevent complement activation, protect mitochondrial integrity, and limit phase separation and drusen nucleation in the RPE.

Our live imaging data provide a basis to understand the reversed risk associations of *APOE* alleles between AMD and Alzheimer’s disease. The dynamic trafficking and efficient cholesterol transport we observed with ApoE4 agrees with studies in astrocytes (31, 66). In the brain, astrocytes deliver cholesterol to neurons in the form of ApoE lipoproteins (67), and high neuronal cholesterol is conducive for the formation of pathogenic beta-amyloid from amyloid precursor protein, suggesting that increased astrocyte-to-neuron cholesterol transport by ApoE4 is injurious to the brain (66). In contrast, in the retina, the RPE is the main biosynthetic source of ApoE, the primary hub for cholesterol trafficking into and out of the retina, and the initiator of drusen biogenesis (6, 8, 35). Cholesterol transport by ApoE occurs in association with ABCA1, and ApoE4 was shown to bind ABCA1 with greater affinity than ApoE3 in CNS astrocytes (ApoE2 was not studied) (68). Genetic variants in ABCA1 are also implicated in AMD (8), and mouse models with RPE-specific deletion of ABCA1 show RPE atrophy and retinal degeneration (17). Thus, in the context of the retina, efficient cholesterol transport mediated by ApoE4 and ABCA1 is likely to be essential for maintaining RPE cholesterol homeostasis and limiting drusen nucleation.

In the RPE, excess cholesterol and ceramide impair intracellular trafficking, leading to autophagic defects and complement-mediated mitochondrial injury (18, 19, 33). These defects were recapitulated in ApoE2-expressing RPE as a direct consequence of poor cholesterol transport and microtubule acetylation. In contrast, efficient cholesterol efflux by ApoE3 and ApoE4 preserved critical functions in RPE exposed to innate stressors (complement, lipofuscin) that are associated with AMD. Autophagic defects and mitochondrial abnormalities have been observed in AMD donor RPE (20, 22, 39, 69), and our studies now provide a molecular mechanism to explain how AMD risk variants in cholesterol genes can drive these phenotypes.

Detailed analyses of ApoE isoform dynamics led to the discovery of an exciting novel mechanism - ApoE liquid-liquid phase separation as an initiator of drusen biogenesis. Proteomics and cell culture studies suggest that drusen originate from the RPE (70). However, these observational studies do not address a critical question in AMD biology: how does ApoE nucleate drusen within the RPE, and can we target this therapeutically? We now show that mitochondrial stress stimulates redox state-mediated phase separation of ApoE2 because of cysteines at positions 112 and 158, whereas ApoE4 (with arginines at both positions) is resistant to phase separation. ApoE3-expressing RPE also undergoes phase separation, albeit to a lesser extent than ApoE2, because of the single cysteine at 112. Given the low prevalence of *APOE2* and *APOE4* alleles in the general populace, these data support the multiple-hit model of AMD pathogenesis where the presence of other genetic (e.g., *CFH, HTRA1/ARMS2, C3)* or environmental (e.g., smoking, high HDL) risks (7, 8, 10, 69) could drive ApoE3 phase separation and drusen nucleation. Indeed, our data on human donors with the *E3/E3* genotype where the extent of RPE mitochondrial injury correlates with the number and volume of ApoE aggregates in AMD donors lend strong support to this model.

Studies in mice confirmed the role of cysteines in mediating protein phase separation: mouse ApoE, which has arginines at 112 and 158 does not form condensates in *Abca4*^-/-^ RPE, despite extensive mitochondrial fragmentation. These data also provide unexpected insight into mouse models of AMD and explain why mice do not develop *bona fide* drusen even in transgenic mice engineered to accumulate cholesterol in the RPE and Bruch’s membrane (71). Our mouse studies led us to another surprising discovery: that mitochondria-mediated phase separation affects not just ApoE2, but other proteins with reactive cysteines, notably p62. Because p62 regulates multiple functions including autophagy, proteasomal clearance, and the antioxidant stress response via NRF2/KEAP1 signaling, phase separation of p62 could compromise many pathways implicated in AMD (62, 72, 73).

Liquid-liquid phase separation of proteins into biomolecular condensates has been increasingly implicated in several neurodegenerative diseases. However, modulating pathological condensate formation requires a deeper understanding of the biology that drives phase transitions (74). Our discovery of mitochondria-mediated phase separation of proteins with reactive cysteines in the RPE, coupled with the relative rarity of cysteines in the proteome (~2.3% (75)), suggests that thiol-mediated selective protein phase separation could be a direct pathological consequence of mitochondrial dysfunction in the RPE. These data also identify a powerful and precise drug target for macular degenerations: rather than simply targeting drusen after they are deposited extracellularly, strategies that act upstream to preserve RPE mitochondrial health could be more beneficial in AMD by halting phase separation, and consequently drusen nucleation within the RPE. In support of this, epidemiological studies show that desipramine - the ceramide-lowering drug used in this study which prevents ApoE2 and p62 phase separation - significantly decreases the incidence of early AMD (76). The RPE is central to understanding disease pathology and developing effective therapies for AMD. These studies identify novel mechanisms that integrate RPE mitochondrial injury with defining, but poorly understood features of AMD: complement activation, lipid dysregulation, and drusen formation. By establishing how ApoE isoforms modulate major AMD-associated biological pathways, our results provide valuable insight into the role of AMD risk variants as “tipping points” that divert the RPE from normal aging towards AMD, and lay the groundwork for further research into the role of liquid-liquid phase separation and biomolecular condensates in AMD pathogenesis.

### Experimental Methods

(Detailed information about reagents, sources, and experimental methods are provided in the Supplementary Information. Brief methods are given below.)

### Primary porcine RPE culture

RPE were isolated from freshly harvested porcine retinas using established protocols (77). For live imaging, primary RPE were plated at confluence (~ 300,000 cells/cm^2^) on serum-coated glass-bottom dishes (Mattek) as described (33, 77).

### Mice

Wild-type (Jackson Labs 129S1/SvImJ) and *Abca4^-/-^* mice (Jackson Labs Abca4tm1Ght/J) both on Rpe65 Leu450 background were raised under 12-h cyclic light with standard diet. Mice (~6-10 months, both sexes) were euthanized ~3 h after light onset. Eyes were enucleated, and eyecups were processed for immunofluorescence staining as detailed previously (18, 19, 33). For desipramine studies, mice were intraperitoneally injected thrice a week for four weeks with either 100 μl sterile, distilled water or 10 mg/kg of desipramine hydrochloride (Enzo Life Sciences, Farmingdale, NY) dissolved in sterile, distilled water (18). All studies were approved by the University of California-San Francisco animal care and use authorities.

### Expression of EGFP or mCherry-tagged ApoE isoforms in primary RPE

pcDNA3.1 plasmids expressing human ApoE2, E3 and E4 were provided by Dr. Joachim Herz (University of Texas Southwestern Medical Center, Dallas, TX) (78). EGFP or mCherry-tagged constructs were generated by inserting the ApoE2, E3 or E4 cDNA into the p-EGFP-N1 or p-mCherry-N1 vector (Invitrogen) between the XhoI and BamHI restriction sites. Sequences of the constructs was confirmed by UW-Madison Biotechnology Core and by Quintara (South San Francisco, CA). Primary porcine RPE were transfected with EGFP or mCherry-tagged ApoE2, E3 or E4 (Amaxa Nucleofector II, Lonza, Rockland, ME). Approximately 1.5 million cells and 5 μg plasmid DNA were used for each transfection. Cells were plated at confluence (300,000 cells/sq.cm) in growth medium on serum-coated glass-bottom dishes (MatTek, Ashland, MA) and used for live imaging 48-72 h later. Transfection efficiencies were ~30-40% for each of the isoforms.

### Treatments and assays

Primary RPE cultures were treated with A2E (10 μM for 6 h and 48 h chase) to match levels found in aging human RPE (33, 79). Other reagents used the ASMase inhibitor desipramine (10 μM, 3 h, Sigma), the LXRa agonist T0901317 (1 μM for 16 h, Cayman Chemicals) and the lipophilic statin Simvastatin (5 μM for 16 h, Cayman Chemicals) (33, 80). Cholesterol levels were measured from RPE lysates using the Amplex Red cholesterol assay kit (ThermoFisher) (33).

### Live imaging and analysis of ApoE trafficking

mCherry-ApoE2, E3 or E4 expressing primary porcine RPE cell cultures were imaged using the Andor Revolution XD with 100X/1.49 NA Apo TIRF objective (Nikon, Melville, NY) for ~50 frames at 37°C. Trafficking data were collected from three separate transfections for a total of at least 17-23 movies captured per condition with the same laser power, exposure and electron-multiplying gain settings for all conditions. For analysis of trafficking parameters, ApoE-labeled vesicles were subjected to surface reconstruction using the Surfaces and Tracks modules and track length, track displacement and track lifetimes were calculated using Imaris v 8.7.4 (Bitplane, Concord, MA).

### Live imaging and analysis of autophagosome biogenesis and trafficking

mCherry-ApoE2, E3 or E4 expressing primary porcine RPE cell cultures were transduced with LC3B-GFP (ThermoFisher, Waltham, MA) for 16-24 hours. Autophagosome trafficking was monitored by live imaging and analysis of autophagosome numbers and trafficking parameters was performed using Imaris as previously described (33).

### Live imaging of mitochondrial dynamics

mCherry-ApoE2, E3 or E4 expressing primary porcine RPE cell cultures ± A2E (10 μM for 6 h followed by a 48-h chase in fresh culture medium) were exposed to 10% NHS for 10 min (19). Cells were incubated with 0.2 μM MitoTracker Deep Red (ThermoFisher, Waltham, MA) for 15 min at 37°C and imaged immediately. For analysis of mitochondrial volume, MitoTracker-labeled mitochondria were subjected to surface reconstruction in Imaris and automated segmentation by color-coding based on volume of the connected components was used for 3D surface rendering of mitochondria.

### Analysis of ApoE condensate dynamics

mCherry-ApoE2, E3 or E4 expressing primary porcine RPE cell cultures were imaged live using the same imaging conditions and parameters established for imaging ApoE trafficking to capture volume, fusion and fission events. In some experiments, mCherry-ApoE2-expressing RPE were treated with 0.5% 1,6 hexanediol (Sigma Aldrich, St. Louis, MO) in digitonin or digitonin alone (vehicle) to disrupt weak hydrophobic interactions or with T0901317, Simvastatin or desipramine as above. Condensate volumes were quantified by 3D surface reconstruction and measuring mCherry fluorescence intensity per pixel of each condensate per z-plane of the confocal image. Histograms of number and volumes of ApoE condensates was constructed using Matlab (MathWorks, Natick, MA).

### Genotyping and immunostaining of human donor globes

Globes from unaffected human donors and donors diagnosed with AMD were obtained from Lions Gift of Sight (Saint Paul, MN) with deidentified demographics (Table S2) and fundus photographs. DNA was extracted from frozen donor retinal tissue (PureLink Genomic DNA Kit, ThermoFisher, Waltham, MA), amplified by PCR using specific primers before genotyping (Quintara) ((8, 22, 81) Table S3). Cryosections of donor retina were stained with antibodies to TOM20 (Santa Cruz sc-11415, 1:200), ApoE (Genetex GTX100053 1:200) or p62/SQSTM1 (Novus NBP148320, 1:200). Sections were labeled with Alexa-Fluor conjugated secondary antibodies, Rhodamine-phalloidin and DAPI to label to label the actin cytoskeleton and nuclei, respectively. Sections were imaged by spinning disc confocal microscopy and analyzed as described below.

### Quantification of mitochondrial volumes and protein condensates

Images of immunostaining were analyzed using Imaris 9.6 (Bitplane). Images were subjected to Gaussian filtering and background subtraction. Surface reconstructions were made with the “Surfaces” module. Mean volumes and areas were exported from the statistics tab and analyzed in Excel and Prism.

### Statistics

Data were analyzed using either one-way ANOVA with Bonferroni post-test or multiple *t*-test with Welch’s correction for unequal variances (GraphPad Prism). Unless otherwise stated, data are presented as mean ± SEM of ≥3 independent experiments, with ≥ three replicates per condition per experiment.

### Study approval

All animal procedures were reviewed and approved by the IACUC at the University of California, San Francisco.

## Supplementary information

Supplementary Information includes detailed Experimental Methods, five figures, eight movies and three tables.

## Acknowledgements

We thank Joachim Herz for the human *APOE2*, *APOE3* and *APOE4* constructs and Jimmy Pham for help with Matlab analyses. This research was supported by NIH/NEI grants R01EY023299 and R01EY030668, the Research to Prevent Blindness/American Macular Degeneration Foundation Catalyst Award for Innovative Research Approaches to AMD, the BrightFocus Foundation Award for AMD research M2015350, Macular Society UK, and the Retina Research Foundation Rebecca Meyer Brown Professorship (AL), NIH/NEI Diversity Supplement Fellowship (NLC) and NIH/NEI P30 core grants for vision research to the University of Wisconsin-Madison (EY016665) and to the University of California, San Francisco (EY002162).

## Author Contributions

NLC, LXT, KAT and AL designed the study; NLC, LXT and KAT performed the experiments; KAT, LXT, GR and CJG generated primary RPE cultures; NLC, GR and CJG processed human donor tissues; TT and NLC performed the genotyping; NLC, LXT, KAT, TT and AL analyzed the data; GR and TT critically reviewed the manuscript; NLC, LX, CJG, KAT and AL wrote the manuscript.

---

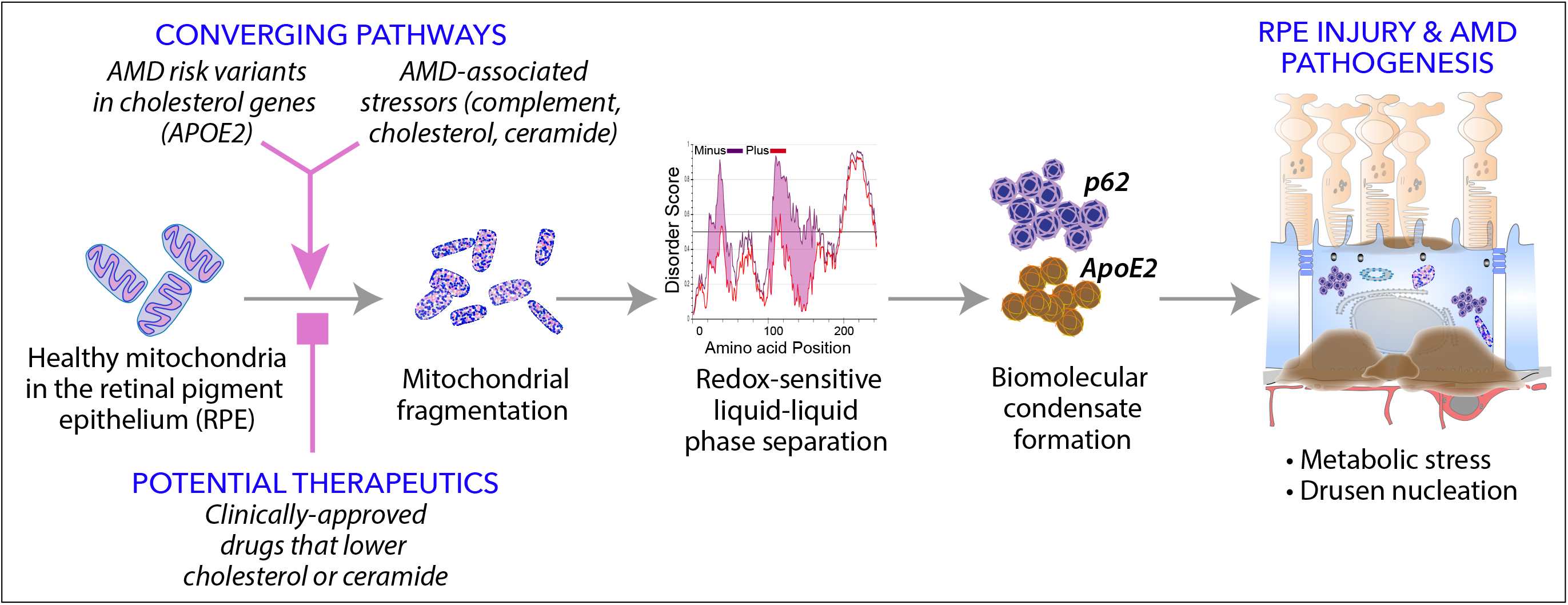

