## Supplemental Info for "Mitochondria-dependent phase separation of disease-relevant proteins drives pathological features of age-related macular degeneration"

**Supplementary Information:**

**Supplementary Figures S1-S5**

**Legends for Supplementary Movies S1 - S8**

**Supplementary Table S1 – Reagents and sources**

**Supplementary Table S2 – Human donor demographics and AMD risk genotypes**

**Supplementary Table S3 – Genotyping information**

**Experimental Methods**

**References**

**A**

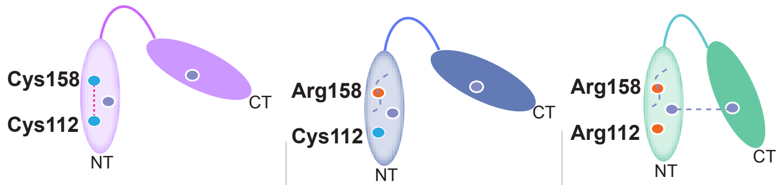

| Feature | ApoE2 | ApoE3 | ApoE4 |
|---|---|---|---|
| Global allele frequency | ~7% | ~79% | ~14% |
| Sequence | Cys112, Cys158 | Cys112, Arg158 | Arg112, Arg158 |
| Tertiary structure | Disruption of N-terminal salt bridge | | Domain interaction (R61-E255 salt bridge) |
| Oligomer formation (intermolecular disulfide bonds) | High propensity | Moderate | Negligible |
| LDLR bindng | <2% | High | High |
| Lipoprotein binding | HDL | HDL | VLDL/HDL |

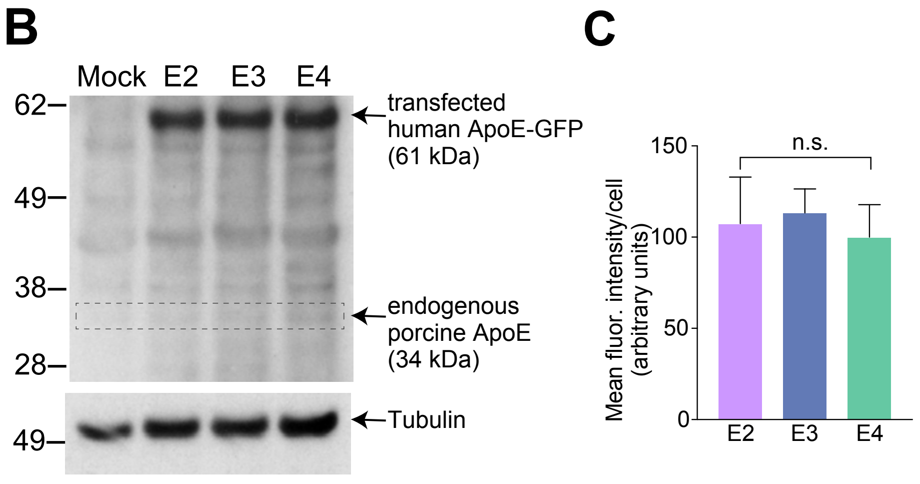

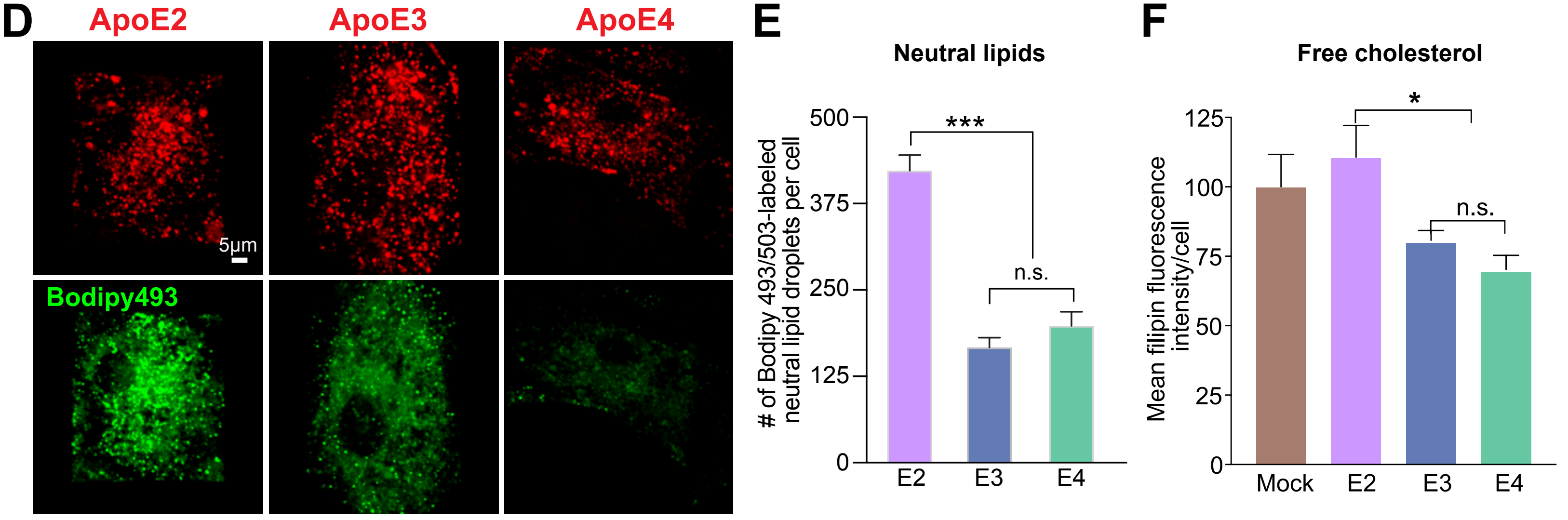

**Figure S1. Human ApoE isoforms - characteristics, expression, and lipid transport. (A)** Structural features and of characteristics of human ApoE isoforms. Key amino acids at positions112 and 158 and potential intermolecular disulfide bonds (red dashed line) and salt bridges (gray dashed lines) are indicated. **(B)** Representative immunoblot of ApoE in porcine primary RPE transfected with EGFP-tagged human ApoE2, E3, or E4. Arrows: 61 kDa fusion proteins, 34 kDa endogenous ApoE and tubulin as loading control. **(C)** Quantification of mean mCherry fluorescence intensity per cell in primary RPE cultures transfected with mCherry-tagged ApoE2, E3 or E4. **(D)** Representative images from live imaging and **(E)** quantification of Bodipy 493/503-labeled neutral lipids in ApoE2, E3 or E4-expressing RPE. Mean ± SEM, n > 36 cells per condition, ***, $p < 0.0001$. **(F)** Primary RPE cultures were stained with filipin to label cholesterol and fluorescence intensity was quantified in cells expressing mCherry--tagged ApoE2, E3 or E4. Mean ± S.E.M. 17-23 cells per condition. One-way ANOVA with Bonferroni post-test*, $p < 0.05$, n.s., not significant.

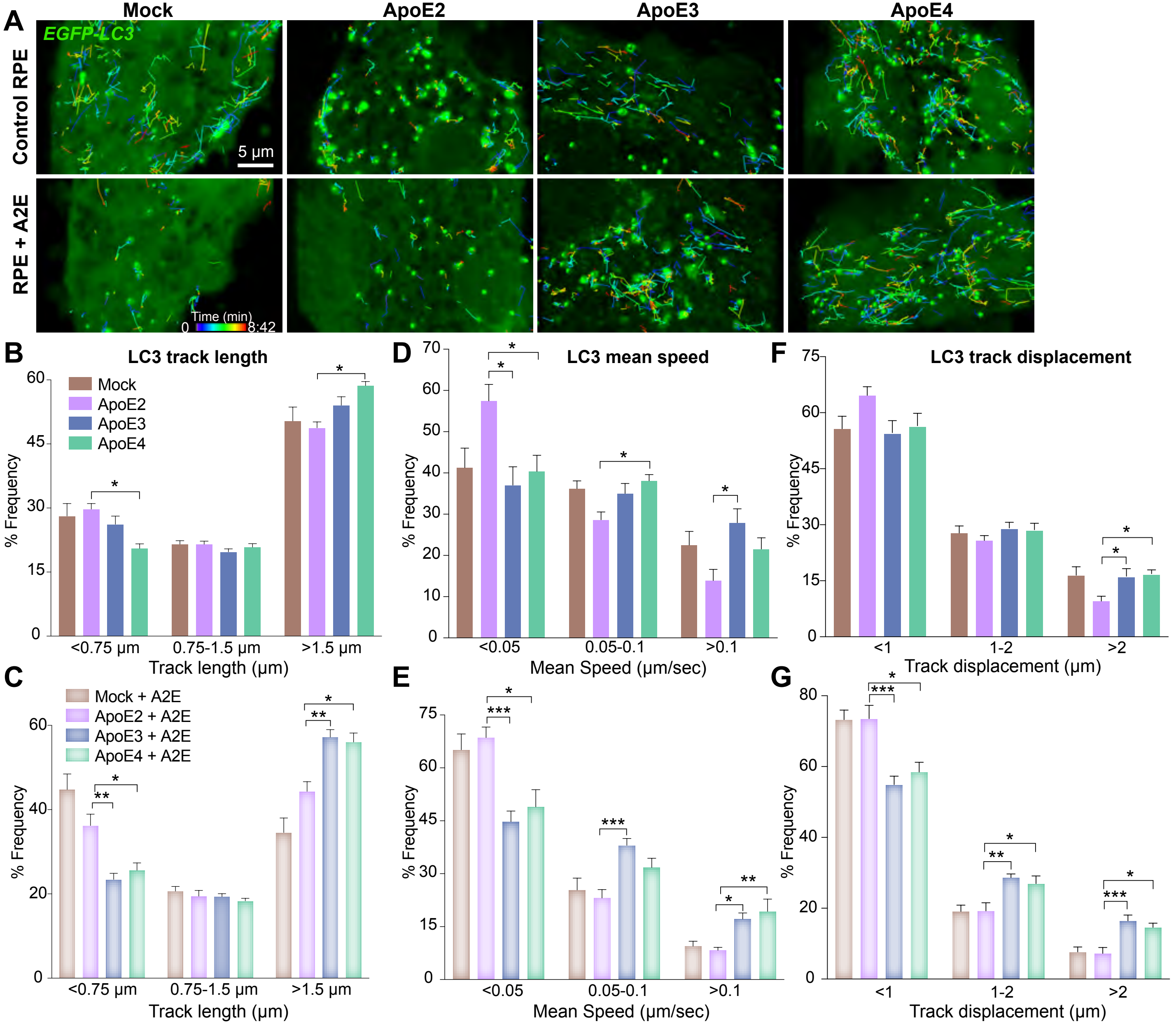

**Supplementary Figure S2. ApoE4 restores autophagosome trafficking in RPE with lipofuscin. (A)** Stills from live imaging of EGFP-LC3 autophagosomes (with spots and tracks superimposed) in mock-transfected or ApoE2, E3 or E4-expressing RPE, and treated or not with A2E. Frequency plots of EGFP-LC3 track lengths **(B)** & **(C)**, mean speeds **(D)** & **(E)**, and track displacements **(F)** & **(G)**. Mean ± SEM, n >13 cells/condition. One-way ANOVA with Bonferroni post-test. *, $p$ < 0.05, **, $p$ < 0.005, ***, and $p$ < 0.001. See also Figure 2 and Movies S1-S8.

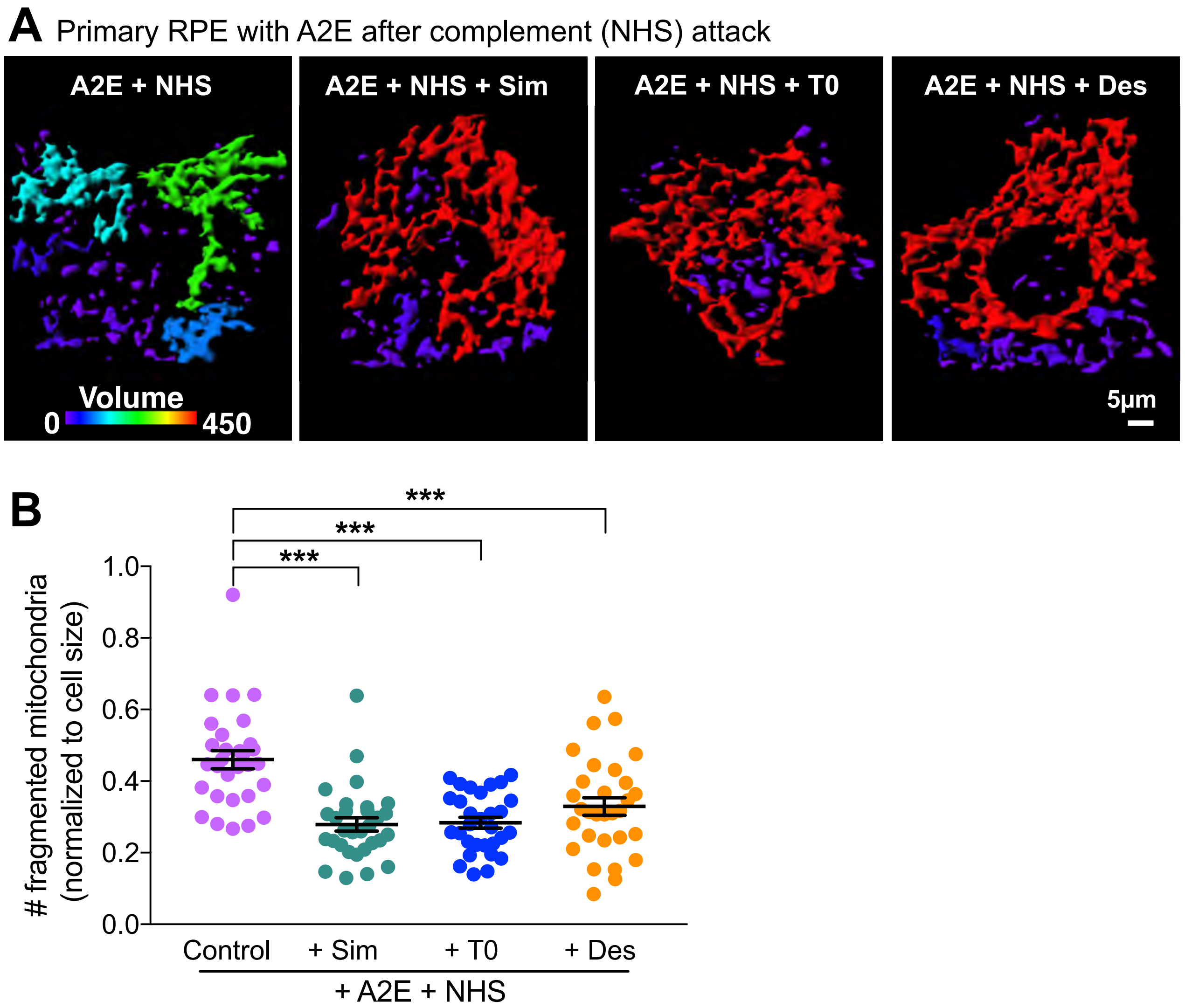

**Figure S3. Drugs that lower cholesterol and ceramide prevent complement-induced mitochondrial fragmentation in RPE with A2E. (A)** 3D reconstruction from live imaging of mitochondrial volumes in mock transfected RPE transduced with Mitotracker, treated with A2E and exposed to 10% NHS to induce complement attack. RPE were treated with Simvastatin (5 μM, 16 h), T0901317 (1 μM, 16 h) or desipramine (10 μM, 3 h) prior to imaging. Color bar: cooler colors indicate increasing mitochondrial fragmentation). **(B)** Number of fragmented mitochondria in (A). ***, $p < 0.0005$. Mean ± SEM, n = 30 cells per condition; One-way ANOVA with Bonferroni post-test; See also Figure 3.

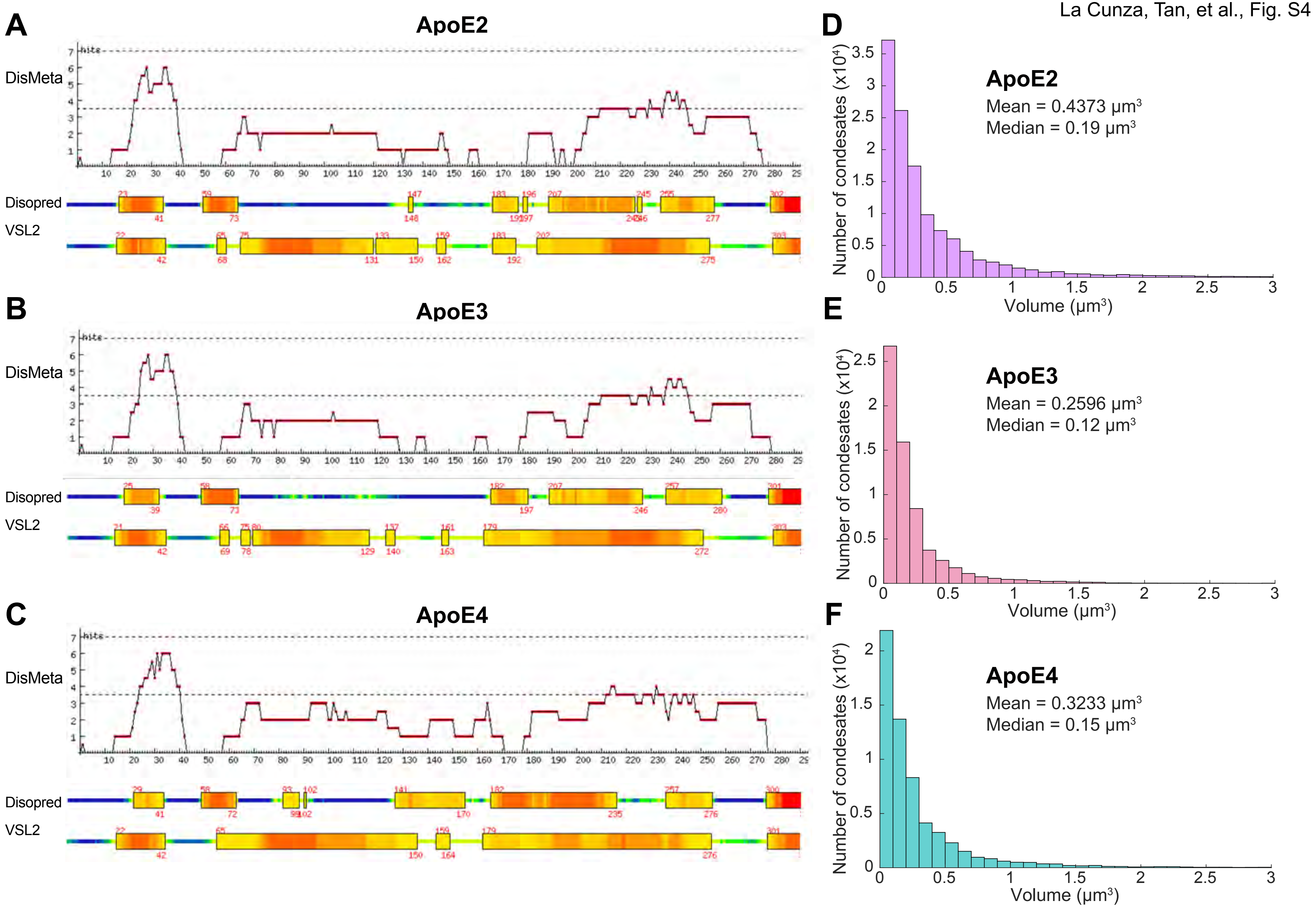

**Figure S4. Phase separaton of ApoE isoforms.** Index of disorder of **(A)** ApoE2, **(B)** ApoE3, and **(C)** ApoE4 predicted by a meta predictor of protein disorder (DisMeta), Disopred and VSL2 (Alberti et al., 2019). **(D), (E), and (F)** Histograms showing number and volumes of ApoE2, E3 or E4 condensates in primary RPE. Note the different scales for the Y-axes.

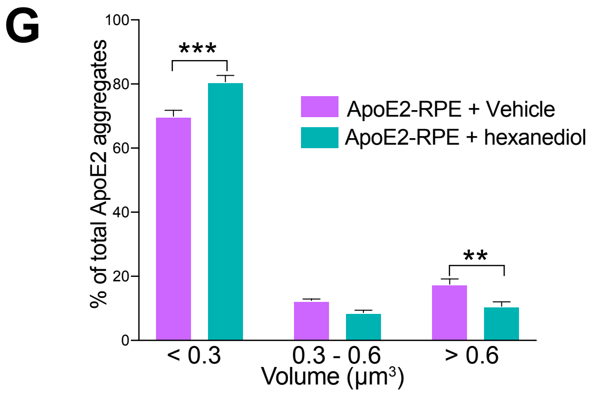

**Figure S4. Redox-sensitive phase separation of ApoE isoforms. (G)** Frequency histogram of ApoE2 condensate volumes treated with digitonin (Vehicle) or 1,6-hexanediol in digitonin. Mean ± SEM, n = 22 cells per condition, **, $p < 0.005$; ***, $p < 0.0001$, one-way ANOVA with Bonferroni post-test.
**(H), (I), (J) and (K)** IUPRED2A disorder plots for human ApoE2, ApoE3, ApoE4 and mouse ApoE. Redox-state dependent order-disorder transitions are depicted by the pink shaded area. Note increased disorder in the region of amino acids 100-200 in ApoE2 **(H)** which spans the two cysteine residues at positions 112 and 158, and between 100-150 in ApoE3 **(I)** that spans the cysteine at position 112. Also note lack of noticeable redox-sensitive order-disorder transition in ApoE4 **(J)** and mouse ApoE **(K)**, which have arginines instead of cysteines at positions 112 and 158.

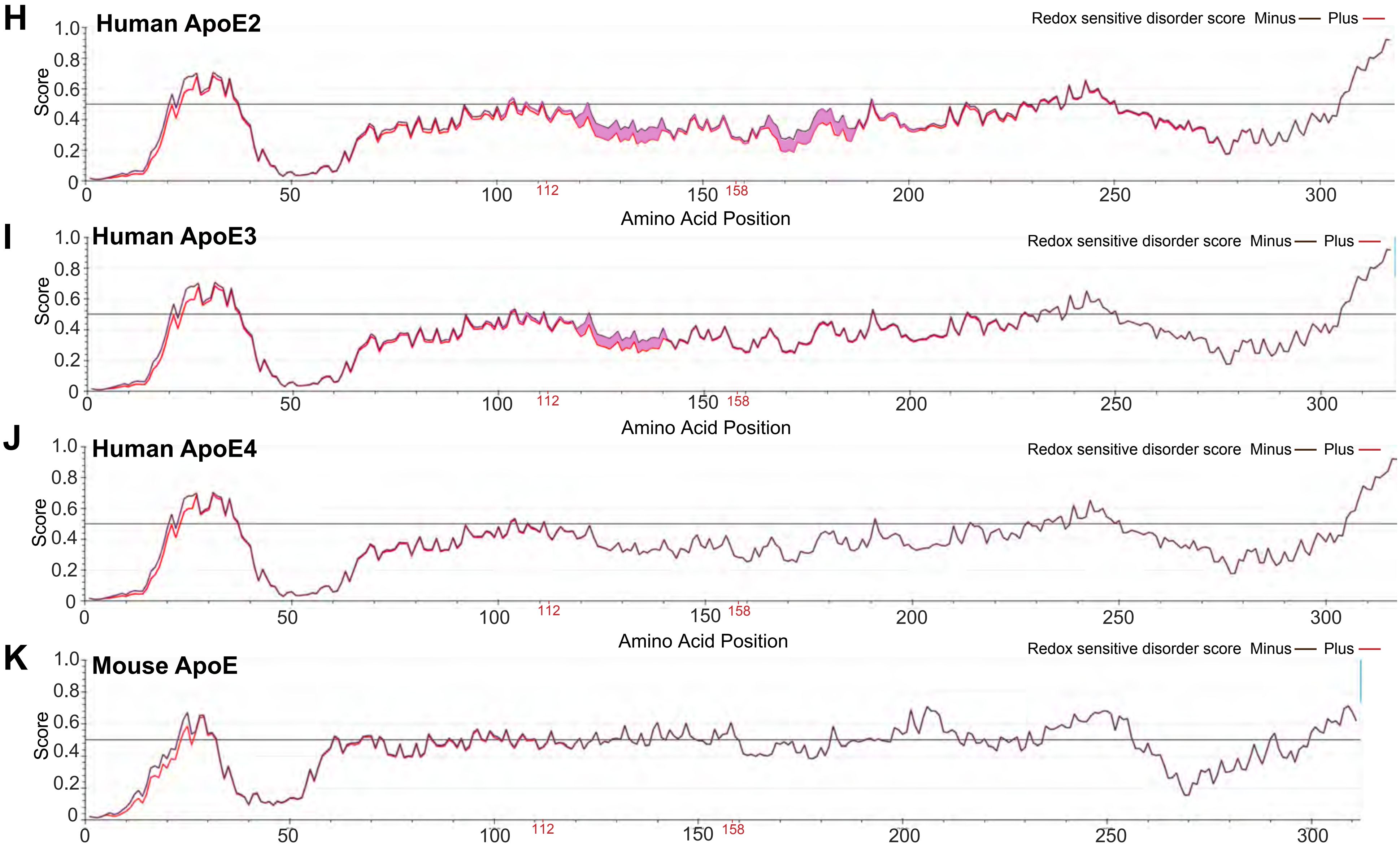

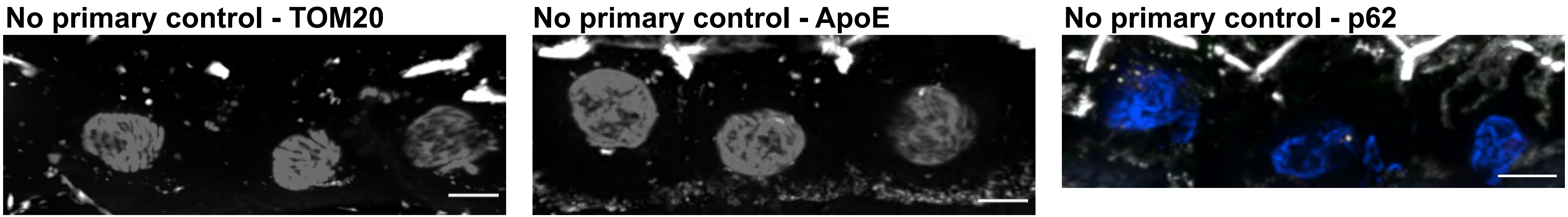

**Figure S5. Antibody controls for immunostaining of human donor cryosections.**
Representative images of immunostaining human donor cryosections incubated with secondary antibodies only. These images were used to control for any fluorescence due to endogenous tissue background (e.g., lipofuscin) and for non-specific binding of the secondary antibody to cellular components. Scale bar = 5 μm

### Legends for Supplementary Movies

**Movie S1:** Live imaging of GFP-LC3 autophagosome trafficking in primary porcine RPE cultures by spinning disk microscopy.

**Movie S2:** Live imaging of GFP-LC3 autophagosome trafficking in A2E-treated primary porcine RPE cultures by spinning disk microscopy.

**Movie S3**. Live imaging of GFP-LC3 autophagosome trafficking in primary porcine RPE cultures expressing mCherry-ApoE2 by spinning disk microscopy.

**Movie S4.** Live imaging of GFP-LC3 autophagosome trafficking in A2E-treated primary porcine RPE cultures expressing mCherry-ApoE2 by spinning disk microscopy.

**Movie S5**. Live imaging of GFP-LC3 autophagosome trafficking in primary porcine RPE cultures expressing mCherry-ApoE3 by spinning disk microscopy.

**Movie S6.** Live imaging of GFP-LC3 autophagosome trafficking in A2E-treated primary porcine RPE cultures expressing mCherry-ApoE3 by spinning disk microscopy.

**Movie S7**. Live imaging of GFP-LC3 autophagosome trafficking in primary porcine RPE cultures expressing mCherry-ApoE4 by spinning disk microscopy.

**Movie S8.** Live imaging of GFP-LC3 autophagosome trafficking in A2E-treated primary porcine RPE cultures expressing mCherry-ApoE4 by spinning disk microscopy.

**Table S1. Reagents and sources**

| REAGENT or RESOURCE | | | SOURCE | IDENTIFIER |
|---|---|---|---|---|
| **Antibodies** | **Host** | **IF or IB conc.** | | |
| Acetylated Tubulin | Mouse | 1:1,000 | Sigma-Aldrich, St. Louis, MO | T6793 |
| ApoE | Rabbit | 1:200 | GeneTex Inc, Irvine, CA | GTX100053 |
| ApoE | Rabbit | 1:300 (IB) | Dako, Glostrup, Denmark | A0077 |
| Beta-tubulin | Mouse | 1:1000 | Sigma, St. Louis, MO | T5201 |
| p62 | Rabbit | 1:200 | Novus Biologicals, Littleton, CO | NBP148320 |
| TOM20 | Rabbit | 1:200 | Santa Cruz Biotechnology, Inc, Dallas, Texas | sc-11415 |
| ZO-1 | Rat | 1:3000 | (Xu et al., 2012 (1)) | |
| Alexa Fluor Secondary antibodies (1:500) – various | | | ThermoFisher Scientific, Waltham, MA | |

| **Plasmids** | |
|---|---|
| pcDNA3.1 plasmids with human ApoE2, ApoE3 or ApoE4 | Dr. Joachim Herz, UT Southwestern (2) |
| mCherry-ApoE2, E3 or E4 | Generated in house from the pcDNA3.1 constructs and sequenced. |
| **Biological Samples** | |
| Human Eyes | Lions Gift of Sight, St Paul, MN |
| **Biological samples: Cells** | |
| Primary porcine RPE | Extracted from freshly harvested pig eyes obtained from Hart & Vold Meat Market, Baraboo, WI and Marin Sun Farms Inc., Petaluma, CA (3) |

| **Biochemicals** | **Source** | **Identifier** |
|---|---|---|
| Bovine Serum Albumin | Rockland Immunochemicals, Pottstown, PA | BSA-50 |
| Calcium Chloride Dihydrate | Sigma-Aldrich, St. Louis, MO | C7902 |
| DAPI (14.3 mM stock solution, used at 1:200) | Sigma-Aldrich, St. Louis, MO | D9542 |
| Digitonin | Sigma Aldrich, St. Louis, MO | D141 |
| dNTP mix | ThermoFisher, Waltham, MA | R72501 |

| | | |
|---|---|---|
| Glucose | Sigma Aldrich, St. Louis, MO | G7528 |
| Glycerol | Sigma Aldrich, St. Louis, MO | G2025 |
| GoTaq G2 | Promega, Madison, WI | M7422 |
| HBSS | Corning, Corning, NY | 21-023-CV |
| HEPES | ThermoFisher, Waltham, MA | 15630080 |
| Magnesium Chloride Hexahydrate | Sigma-Aldrich, St. Louis, MO | M2393 |
| NaCl | Sigma-Aldrich, St. Louis, MO | S5886 |
| Paraformaldehyde (8%) | Electron Microscopy Sciences, Hatfield, PA | 157-8 |
| Phosphate Buffer Saline | ThermoFisher Scientific, Waltham, MA | BP665-1 |
| Protease Inhibitors (used at 1:200) | Calbiochem, San Diego, CA | 539134 |
| Saponin | Sigma-Aldrich, St. Louis, MO | 84510 |
| Sucrose | Sigma-Aldrich, St. Louis, MO | S0389 |
| Triton X-100 | Sigma-Aldrich, St. Louis, MO | T8532 |
| Trizma Base | Sigma-Aldrich, St. Louis, MO | RDD008 |
| TrueBlack | Biotium, Fremont, CA | 23007 |
| Tween-20 | Sigma-Aldrich, St. Louis, MO | P9416 |
| VectaShield | Vector Laboratories, Burlingame, CA | H1000 |

| **Biochemicals: Components for porcine RPE cell culture** | | |
|---|---|---|
| Ciprofloxacin | Sigma-Aldrich, St. Louis, MO | 17850-5G-F |
| DMEM | Corning, Corning, NY | 10-013-CV |
| Fetal Bovine Serum (heat-inactivated) | American Type Culture Collection, Manassas, VA | 30-2020 |
| Non-essential amino acids (NEAA) | Corning, Corning, NY | 25-025-CI |
| Penicillin-Streptomycin | Corning, Corning, NY | 30-002-CI |
| 0.25% Trypsin | Corning, Corning, NY | 25-053-CI |
| 2.5% Trypsin | Lonza, Walkersville, MD | 17-160E |

| **Biochemicals: Dyes** |
|---|

| | | |
|---|---|---|
| Bodipy 493/503 | ThermoFisher, Waltham, MA | D3922 |
| Filipin III | Sigma-Aldrich, St. Louis, MO | F4767 |
| Mitotracker Far Red | ThermoFisher, Waltham, MA | M22426 |
| Rhodamine phalloidin | Cytoskeleton Inc., Denver, CO | PHDR1 |

| **Biochemicals: Pharmacological treatments** | | |
|---|---|---|
| A2E | Synthesized and purified according to published protocols (4-6). | |
| Simvastatin | Cayman Chemicals, Ann Arbor, MI, USA | 10010344 |
| T0901317 | Cayman Chemicals, Ann Arbor, MI, USA | 71810 |
| Desipramine Hydrochloride | Enzo Life Sciences, Farmingdale, NY | BML-AR119-0100 |
| 1,6 Hexanediol | Sigma Aldrich, St. Louis, MO | 240117 |
| Normal Human Serum | Quidel Corporation, San Diego, CA | A113 |

| **Critical Commercial Assays** | | |
|---|---|---|
| Amplex Red Cholesterol Assay Kit | ThermoFisher, Waltham, MA | A12216 |
| DC protein assay kit | Bio-Rad Laboratories, Hercules, CA | 5000112 |
| PureLink® Genomic DNA Kit | ThermoFisher, Waltham, MA | K182001 |
| Phusion™ High-Fidelity DNA Polymerase | ThermoFisher, Waltham, MA | F530S |

| **Materials** | | |
|---|---|---|
| 4-12% NuPAGE® Bis-Tris gels | Invitrogen, Carlsbad, CA | NP0321BOX |
| Embedding molds | ThermoFisher, Waltham, MA | 50-465-348 |
| T25 culture flasks | Corning, Corning, NY | 353108 |
| MatTek dish (35mm) | MatTek Corporation, Ashland, MA | P35G-1.5-14-G |
| TissueTek OCT | Radnor Corporate Center, Radnor, PA | 25608-930 |

| **Transduction reagents** |
|---|

| | | |
|---|---|---|
| Premo Autophagy LC3B-GFP | ThermoFisher, Waltham, MA | P36235 |

| **Software and Algorithms** | |
|---|---|
| GraphPad Prism 8 | GraphPad Prism®, La Jolla, CA |
| Image Studio Lite Ver. 5.2.5 | LI-COR, Lincoln, NE |
| Imaris v 8.7.4 | Bitplane, Concord, MA |
| Matlab R2020a | MathWorks, Natick, MA |

**Table S2. Demographic and genotyping information of human donors**
(Risk allele is in red)

| *Donor #* | **Donor Id#** | **Donor demographics** | **Clinical diagnosis** | ***APOE*** | ***CFH*** | ***C3*** | ***HTRA1*** | ***ARMS2*** | **Cause of death** | **Death to preservation** |
|---|---|---|---|---|---|---|---|---|---|---|
| *1* | 996 | 65 (Male, Caucasian) | Control | ***E3/E4*** *rs429358 C/T* *rs7412 C/C* | CT | CG | GG | GG | Lung cancer | 10 hours |
| *2* | 226 | 76 (Male, Caucasian) | Control | ***E3/E3*** *rs429358 T/T* *rs7412 C/C* | TT | CG | AG | GT | Respiratory failure, pneumonia | 13 hours |
| *3* | 684 | 94 (Female, Caucasian) | Control | ***E3/E3*** *rs429358 T/T* *rs7412 C/C* | TT | CG | GG | GG | Colon cancer | 7 hours |
| *4* | 236 | 95 (Female, Caucasian) | AMD | ***E3/E3*** *rs429358 T/T* *rs7412 C/C* | CT | CG | GG | GG | Small bowel obstruction | 6 hours |
| *5* | 1004 | 84 (Male, Caucasian) | AMD | ***E3/E3*** *rs429358 T/T* *rs7412 C/C* | CT | CC | GG | GT | Respiratory failure | 6 hours |
| *6* | 1061 | 82 (Male, Caucasian) | AMD | ***E3/E4*** *rs429358 C/T* *rs7412 C/C* | TT | CG | AG | GT | Abdominal aortic aneurysm | 6 hours |

**Table S3. PCR primers and conditions for genotyping**

| Gene | SNP | Primer | PCR Conditions | Size (bp) |
|---|---|---|---|---|
| CFH Y402H | rs1061170 | F: 5'- TCATTGTTATGGTCCTTAGGAAA-3'<br>R: 5'- TTAGAAAGACATGAACATGCTAGG-3' | 98ºC for 3 min,<br>98ºC for 10 sec,<br>58ºC for 45 sec,<br>72ºC for 45 sec,<br>72ºC for 7 min;<br>34 cycles | 241 |
| C3<br>R102G | rs2230199 | F: 5'- GACAAAGAGGCCTCGTGAGA-3'<br>R: 5'- GACCAAGAATAATGGGCAGGC-3' | 98ºC for 3 min,<br>98ºC for 10 sec, 64ºC for 20 sec, 72ºC for 20 sec, 72ºC for 5 min;<br>34 cycles | 348 |
| ARMS2 | rs10490924 | F: 5'- TACCCAGGACCGATGGTAAC-3'<br>R: 5'-GAGGAAGGCTGAATTGCCTA-3 | 98ºC for 3 min,<br>98ºC for 10 sec, 58ºC for 20 sec, 72ºC for 40 sec, 72ºC for 7 min;<br>35 cycles | 449 |
| HTRA1 | rs11200638 | F: 5'- CTAGGCTCTCTGCGAATACGG-3'<br>R: 5'- ATGGAACTTTGCAAGGGGGC-3' | 98ºC for 3 min,<br>98ºC for 10 sec, 64ºC for 20 sec, 72ºC for 20 sec, 72ºC for 5 min;<br>34 cycles | 396 |
| APOE | rs429358 &<br>rs7412 | F: 5'-TCGGAACTGGAGGAACAACT-3'<br>R: 5'-ACCTGCTCCTTCACCTCGT-3' | 98ºC for 3 min,<br>98ºC for 10 sec, 64ºC for 20 sec, 72ºC for 20 sec, 72ºC for 5 min;<br>34 cycles | 482 |

### Experimental Methods

#### Primary porcine RPE cell cultures

RPE were isolated from porcine retinas using established protocols (3). Briefly, the anterior segment was removed at the *ora serrata*, and the retina was gently detached by clipping at the optic nerve head. RPE cells were isolated from eyecups upon incubation with 0.5% trypsin (Lonza, Walkersville, MD) with 5.3 mM EDTA in Hank's balanced salt solution (Corning, Corning, NY) and plated onto T25 flasks in growth medium (DMEM with 4.5 g/L glucose, 1% heat-inactivated fetal bovine serum, 1% non-essential amino acids and 1% penicillin/streptomycin). For live imaging, RPE were plated at confluence (~ 300,000 cells/cm$^2$) on serum-coated glass-bottom dishes (Mattek) as described (3, 5).

#### Treatments and assays

Pharmacological treatments were performed as previously described (3, 7, 8). Cells were treated with 10 µM bisretinoid A2E for 6 h followed by a 48-h chase in fresh culture medium. The LXR agonist T0901317 (Cayman Chemicals, Ann Arbor, MI) was used at 1 µM for 16 h, the ASMase inhibitor desipramine hydrochloride was used at 10 µM for 3 h and the lipophilic statin simvastatin (Cayman Chemicals, Ann Arbor, MI) was used at 5 µM for 16 h. Total cell cholesterol levels were measured with the Amplex Red cholesterol assay (ThermoFisher) as detailed previously (5, 6).

#### ApoE constructs

pcDNA3.1 plasmids expressing human ApoE2, E3 and E4 were provided by Dr. Joachim Herz (University of Texas Southwestern Medical Center, Dallas, TX) (2). The mCherry-tagged constructs were generated by inserting the ApoE2, E3 or E4 cDNA into the p-mCherry-N1 vector (Invitrogen) between the XhoI and BamHI restriction sites. Sequences of the constructs was confirmed by UW-Madison Biotechnology Core and by Quintara (South San Francisco, CA).

#### Transfections

Primary porcine RPE were transfected with mCherry-tagged ApoE2, E3 or E4 using the Amaxa nucleofector II (Lonza, Rockland, ME). Approximately 1.5 million cells and 5 µg plasmid DNA were used for each transfection. Cells were plated at confluence (300,000 cells/sq.cm) in growth medium on serum-coated glass-bottom dishes (MatTek, Ashland, MA) and used for live imaging 48-72 h later. Transfection efficiencies were ~30-40% for each of the isoforms.

**Live imaging and analysis of ApoE trafficking**

mCherry-ApoE2, E3 or E4 expressing primary porcine RPE cell cultures were imaged in recording medium (1 x HBSS, 4.5 g/L glucose, 0.01 M HEPES) supplemented with 1% FBS and 1% NEAA as described previously (2,4,5). Rapid Z-stacks were acquired using the Andor Revolution XD with 100x/1.49 NA Apo TIRF objective (Nikon, Melville, NY) for ~50 frames at 37°C. Trafficking data were collected from three separate transfections for a total of at least 17–23 movies captured per condition. During image acquisition, care was taken to maintain the same laser power, exposure and electron-multiplying gain settings. Images were subjected to background subtraction and Gaussian smoothing (same settings for all images within a set of experiments) in Imaris v 8.7.4 (Bitplane, Concord, MA). For analysis of trafficking parameters, ApoE-labeled vesicles were subjected to surface reconstruction using the Surfaces and Tracks modules, with the same intensity threshold applied to all images within a set of experiments. Track length, track displacement and track lifetimes were exported and statistical analysis of these data sets was performed using Prism (GraphPad, La Jolla, CA).

**Immunoblotting**

For ApoE immunoblotting, primary porcine RPE pellets were lysed in 1X HNTG lysis buffer (50mM HEPES, 150mM NaCl, 10% glycerol, 1.5mM MgCl2, 1% Triton X-100) supplemented with protease inhibitors (1:200, Calbiochem, San Diego, CA, 539134) by sonication for 10 min. Protein measurements were performed using DC assay kit (Bio-Rad Laboratories, Hercules, CA). 30 μg of lysates per condition were resolved in 4-12% NuPAGE® Bis-Tris Precast gels (Invitrogen, Carlsbad, CA) at 130 V for ~1.5 h, transferred onto nitrocellulose membrane with iBlot dry transfer system (Thermo Fisher, Waltham, MA). Membranes were blocked with 5% milk in Tris buffered saline with Tween20 (TBS-T) for 1 h before being incubated in primary antibody overnight at 4ºC. Membrane was probed with antibodies to rabbit anti-human ApoE (1:300; Dako, #A0077, Glostrup, Denmark), mouse-anti beta-tubulin (1:1000; Sigma #T5201, St. Louis, MO) followed by washes in TBS-T and incubated with horseradish peroxidase-conjugated secondary antibodies (1:5000) for 1 h. After immunoblots were visualized by ECL substrate (Thermo

Fisher, Waltham, MA), bands were quantified using Image Studio (LI-COR, Lincoln, NE) and ApoE levels were normalized to that of beta-tubulin.

**Neutral lipid staining with Bodipy 493/503**

mCherry-ApoE2, E3 or E4 expressing primary porcine RPE cell cultures were treated with Bodipy-493/503 (10 µM for 30 min at 37ºC). Cells were washed, fixed in 2% paraformaldehyde and imaged using the Andor Revolution XD with 100x/1.49 NA Apo TIRF objective (Nikon, Melville, NY). The number of Bodipy493/503 positive vesicles were quantified by the Spots module (Imaris). Statistical analysis of these data sets was performed using Prism (GraphPad, La Jolla, CA).

**Free cholesterol staining with filipin**

mCherry-ApoE2, E3 or E4 expressing primary porcine RPE cell cultures were fixed with 2% paraformaldehyde and stained with 50 µg/ml filipin for 45 min (F4767, Sigma-Aldrich, St. Louis, MO) and imaged using the Andor Revolution XD with 100x/1.49 NA Apo TIRF objective (Nikon, Melville, NY). Intracellular filipin fluorescence was quantified by using Imaris. Statistical analysis of these data sets was performed using Prism (GraphPad, La Jolla, CA).

**Live imaging of autophagosome biogenesis and trafficking**

mCherry-ApoE2, E3 or E4 expressing primary porcine RPE cell cultures were transduced at 6.7 transducing units/cell of Premo Autophagy LC3B-GFP (ThermoFisher, Waltham, MA) for 16-24 hours. Cells were serum starved 2 h to induce autophagy and imaged in recording medium (1 x HBSS, 4.5 g/L glucose, 0.01 M HEPES) supplemented with 1% FBS and 1% NEAA as described previously (5). Time-lapse movies were acquired using the Andor Revolution XD with 100x/1.49 NA Apo TIRF objective (Nikon, Melville, NY) for ~50 frames (50 millisecond exposure) at 37°C. Trafficking data were collected from three separate transfections for a total of at least 12–15 movies captured for treatment. Identical laser power, exposure and gain settings were applied within the same set of experiments. Images were subjected to background subtraction and Gaussian smoothing (same settings for all images within a set of experiments) in Imaris v 8.7.4 (Bitplane, Concord, MA). For analysis of organelle numbers and trafficking parameters, LC3B-labeled autophagosomes were subjected to spots reconstruction using the Spots and Tracks modules, with the same intensity threshold applied to all images within a set of experiments. Numbers, track length, track mean speed and track displacement length were exported and statistical analysis of these data sets was performed using Prism (GraphPad, La Jolla, CA) as described previously (5).

**Acetylated tubulin staining and quantification**

mCherry-ApoE2, E3 or E4 expressing primary porcine RPE cell cultures were treated or not with A2E (10 µM for 6 h followed by a 48-h chase in fresh culture medium). Cells were fixed in 2% paraformaldehyde for 10 min at RT, rinsed in PBS with $Ca^{2+}$ and $Mg^{2+}$ (1x PBS with 1M $MgCl_2$ and 1M $CaCl_2$; PBS C/M), blocked in 1% bovine serum albumin (BSA) in PBS C/M, and incubated with specific primary antibodies diluted in 1% BSA in PBS C/M and 1% saponin for 1 h: mouse monoclonal anti-acetylated tubulin clone 6-11B-1 (1:1000; Sigma-Aldrich), and rat anti-ZO1 (1:3000; Xu et al., 2012). Alexa Fluor secondary antibodies were used at 1:500 and DAPI (1:200; Sigma-Aldrich, St. Louis, MO). Cells were imaged with an Andor Revolution XD with 100x/1.49 NA Apo TIRF objective (Nikon, Melville, NY) with identical exposures and gains for each antibody. For each transfected cell, a region of interest (ROI) was drawn just within the plasma membrane, and the acetylated tubulin fluorescence intensity per pixel of the ROI per plane of the confocal image stack was quantified by Imaris. Statistical analysis of these data sets was performed using Prism (GraphPad, La Jolla, CA).

**Live imaging of mitochondrial dynamics**

mCherry-ApoE2, E3 or E4 expressing primary porcine RPE cell cultures ± A2E (10 µM for 6 h followed by a 48-h chase in fresh culture medium) were exposed to 10% NHS for 10 min (on day 1 and day 3, with a rest on day 2 (8)). Cells then incubated with 0.2 µM MitoTracker Deep Red (ThermoFisher, Waltham, MA) for 15 min at 37ºC and imaged immediately. Some cultures were treated with the LXR agonist T0901317 (Cayman Chemicals, Ann Arbor, MI) at 1 µM for 16 h or the ASMase inhibitor desipramine hydrochloride at 10 µM for 3 h or the lipophilic statin simvastatin (Cayman Chemicals, Ann Arbor, MI) at 5 µM for 16 h. RPE cells were incubated with 0.2 µM MitoTracker Deep Red (ThermoFisher, Waltham, MA) for 15 min at 37ºC immediately prior to imaging. Cells were imaged in recording medium (1 x HBSS, 4.5 g/L glucose, 0.01 M HEPES) supplemented with 1% FBS and 1% NEAA as described previously (8) on the Andor

Revolution XD with 100x/1.49 NA Apo TIRF objective (Nikon, Melville, NY) with 50 millisecond exposure at 37°C. Identical laser power, exposure and gain settings were applied within the same set of experiments. Images were subjected to background subtraction and Gaussian smoothing (same settings for all images within a set of experiments) in Imaris v 8.7.4 (Bitplane, Concord, MA). For analysis of mitochondrial volume, MitoTracker-labeled mitochondria were subjected to surface reconstruction using the Surfaces and Tracks modules, with the same intensity threshold applied to all images within a set of experiments. The Surface module takes a 3D data set, as from confocal imaging, and uses an object identification algorithm with user defined threshold criteria to build artificial solid objects within the image that can be measured for statistics such as the number of discrete or connected objects and the object's volumes. Automated segmentation of these objects by color-coding, based on volume of the connected components, was used for 3D surface rendering of these objects. Statistical analysis of these data sets was performed using Prism (GraphPad, La Jolla, CA).

**Analysis of ApoE condensate volumes**

For analysis of ApoE condensate volumes, mCherry-ApoE2, E3 or E4 expressing primary porcine RPE cell cultures were imaged live using the same imaging conditions and parameters established for imaging ApoE trafficking. The mCherry-ApoE structures were subjected to surface reconstruction using the Surfaces and Tracks modules, with the same intensity threshold applied to all images within a set of experiments. For each transfected cell, a region of interest (ROI) was drawn and the mCherry fluorescence intensity per pixel of the ROI per plane of the confocal image stack was quantified by Imaris. Volumes were exported from the Statistics tab. Histograms of number and volumes of ApoE condensates were created by Matlab (MathWorks, Natick, MA). Statistical analysis of these data sets was performed using Prism (GraphPad, La Jolla, CA).

**Analysis of ApoE condensates after 1,6-hexanediol treatment**

mCherry-ApoE2 expressing primary porcine RPE cell cultures were treated with 5 $\mu$g/mL digitonin alone (control) for 15 min or 5 $\mu$g/mL digitonin (Sigma Aldrich, St. Louis, MO) for 13 min with the aliphatic alcohol, 0.5% 1,6 hexanediol (Sigma Aldrich, St. Louis, MO) added for the last 2 min of the 15 min treatment (9). Cells were imaged in recording medium (1 x HBSS, 4.5 g/L glucose, 0.01 M HEPES) supplemented with 1% FBS and 1% NEAA using spinning disk confocal microscopy (Nikon CSU-X1 dual camera platform equipped with Okolab stagetop incubation system: specifically, the iXon Ultra 888 EM-CCD camera) using a 100X Apo TIRF 1.49 NA oil objective for ~25 frames (50 millisecond exposure) at 37ºC. Identical laser power, exposure and gain settings were applied within the same set of experiments. Images were subjected to background subtraction and Gaussian smoothing (same settings for all images within a set of experiments) in Imaris v 8.7.4 (Bitplane, Concord, MA). For analysis of numbers and volume, ApoE condensates were subjected to surface reconstruction using the Surfaces and Tracks modules, with the same intensity threshold applied to all images within a set of experiments. Histogram of number and volumes of ApoE2 condensates was created by Matlab (MathWorks, Natick, MA). Statistical analysis of these data sets was performed using Prism (GraphPad, La Jolla, CA).

**Live imaging of ApoE condensate fusion**

mCherry-ApoE2, E3 or E4 expressing primary porcine RPE cell cultures were imaged on a spinning disk confocal microscope (Nikon CSU-X1 dual camera platform equipped with Okolab stagetop incubation system: specifically, the iXon Ultra 888 EM-CCD camera) using a 100X Apo TIRF 1.49 NA oil objective at 37ºC. To capture ApoE condensate fusion events, time-lapse movies were acquired at 9 s intervals for 24 seconds (50 millisecond exposure) at 37°C. Identical laser power, exposure and gain settings were applied within the same set of experiments. Images were subjected to background subtraction and Gaussian smoothing (same settings for all images within a set of experiments) in Imaris v 8.7.4 (Bitplane, Concord, MA).

**Mice**

Wild-type (Jackson Labs 129S1/SvImJ) and *Abca4*$^{-/-}$ mice (Jackson Labs Abca4tm1Ght/J) both on Rpe65 Leu450 background were raised under 12-h cyclic light with standard diet. Mice (~6-10 months, both sexes) were euthanized ~3 h after light onset. Eyes were enucleated, and eyecups were processed for immunofluorescence staining as detailed previously (5, 7, 8). For desipramine studies, mice were intraperitoneally injected thrice a week for four weeks with either 100 µl sterile, distilled water or 10 mg/kg of desipramine hydrochloride (Enzo Life Sciences, Farmingdale, NY) dissolved in sterile, distilled water (7). All studies were approved by the University of California-San Francisco animal care and use authorities.

**Human donor eyes**

Globes from unaffected human donors and donors diagnosed with AMD were obtained from Lions Gift of Sight (Saint Paul, MN) with de-identified demographics (Table S2) and fundus photographs. Globes were enucleated within 6-10 hours after death, fixed in 4% PFA and shipped in normal saline. The anterior portions of the globe were removed by cutting along the *ora serrata* and the cornea, lens and vitreous were discarded. The eyecups were infiltrated with 10%, 20%, 30% sucrose in phosphate buffer for 30 min each, followed by 4:1 30% sucrose:OCT and 2:1 30% sucrose:OCT for 30 min each. The macula and periphery regions of the RPE were separated, frozen in TissueTek OCT in embedding molds and sectioned into 20 µm cryosections using a Leica microtome.

**Genotyping of human donor tissue**

DNA samples from flash frozen human eye globes (Lions Gift of Sight) were extracted from ~20 mg eye tissue by using the PureLink Genomic DNA Kit (ThermoFisher, Waltham, MA) according to the manufacturer's instructions. DNA was diluted with nuclease free water to 40 ng/µL for genotyping analysis. PCR amplification was performed as follows for each SNP. For ABCA1 and CETP, reactions were carried out in a total volume of 20µL containing 40-50 ng DNA, 10pM of each primer, GoTaq G2 buffer, and nuclease free water. For CFH Y402H, reactions were carried out in a total volume of 20 µL containing 40 ng DNA, 2.5 mM dNTP mix, 10 µM of each primer, 5x Phusion GC Buffer, Phusion DNA polymerase (2 U/µL) and nuclease free water. For the other variants, reactions were carried out in a total volume of 20 µL containing 40 ng DNA, 2.5 mM dNTP mix, 10 µM of each primer, 5x Phusion GC Buffer, 100% DMSO, Phusion DNA polymerase (2 U/µL) and nuclease free water. Phusion™ High-Fidelity DNA Polymerase kit (ThermoFisher, Waltham, MA) was used according to the manufacturer's instructions. The PCR primers and conditions are summarized in Table S3 (10-12). All PCR products were sent to Quintara (South San Francisco, CA) for Sanger sequencing and the genotyping results are in Table S2.

**Immunofluorescence staining of human and mouse cryosections**

Cryosections were thawed for 10 min at room temperature. Cryosections were blocked in 4% BSA (Rockland Immunochemicals, Pottstown, PA) in PBS (ThermoFisher, Waltham, MA) for 2.5 hrs at room temperature. After three washes in 1% BSA in PBS, cryosections were incubated with the following antibodies in 1% BSA in PBS at 4ºC for 48 h in a humidified chamber (rabbit anti-ApoE (Genetex; 1:200), rabbit anti-p62/SQSTM1 (Novus; 1:200) and rabbit anti-TOM20 (Santa Cruz; 1:200)). Sections were washed three times with PBS to remove unbound antibodies, and incubated with AlexaFluor secondary antibodies (1:500, ThermoFisher Scientific, Waltham, MA) (16 h, 4ºC for human and 2 h at room temperature for mouse cryosections). After one wash in PBS with 1% BSA, the sections were incubated with Rhodamine phalloidin (14 µM; 1:200, Cytoskeleton Inc, Denver, CO, PHDR1) (30 min, RT) and DAPI (14.3 mM; 1:200, Sigma-Aldrich, St. Louis, MO, D9542) (10 min, RT). Human cryosections were treated with TrueBlack reagent (Biotium, Fremont, CA, 23007) for 30 s after immunostaining and rinsed with PBS (3 times, 5 mins each) to quench lipofuscin autofluorescence. Cryosections were mounted under glass coverslips with PBS and Vectashield (Vector Laboratories, Burlingame, CA). Sections were imaged on a spinning disk confocal microscope (Nikon CSU-X1 dual camera platform using the Photometrics Prime 95B Back-illuminated sCMOS) using a 100X Apo TIRF 1.49 NA oil objective with identical exposures and gains for each antibody.

**Analysis of mitochondrial volumes and protein condensates**

Human donor cryosections and mouse RPE flatmounts were immunostained with antibodies to Tom20 (1:200, Santa Cruz), ApoE (Genetex; 1:200) or p62 (1:200, Novus) and analyzed with Imaris 9.6 (Bitplane). Images were subjected to Gaussian filtering and background subtraction. Surface reconstructions were made with the "Surfaces" module. Mean volumes and areas were exported from the statistics tab and analyzed in Excel and Prism.

**Statistics**

Data were analyzed by using either a one-way or two-way ANOVA with the Bonferroni post hoc test or a multiple *t*-test with Welch's correction for unequal variances (GraphPad Prism). Unless otherwise stated, data are presented as mean ± SEM of ≥3 independent experiments, with at least two to three replicates per condition per experiment.
